# Identification of a VPS29 isoform with restricted association to Retriever and Retromer accessory proteins through auto-inhibition

**DOI:** 10.1101/2025.02.04.636264

**Authors:** James L. Daly, Kai-en Chen, Rebeka Butkovič, Qian Guo, Michael D. Healy, Eva Pennink, Georgia Gamble-Strutt, Zara Higham, Edmund R.R. Moody, Philip A. Lewis, Kate J. Heesom, Tom A. Williams, Kirsty J. McMillan, Brett M. Collins, Peter J. Cullen

## Abstract

The endosomal-lysosomal network is a hub of organelles that orchestrate the dynamic sorting of hundreds of integral membrane proteins to maintain cellular homeostasis. VPS29 is a central conductor of this network through its assembly into Retromer, Retriever and Commander endosomal sorting complexes, and its role in regulating RAB GTPase activity. Two VPS29 isoforms have been described, VPS29A and VPS29B, that differ solely in their amino-terminal sequences. Here we identify a third VPS29 isoform, which we term VPS29C, that harbours an extended amino-terminal sequence compared to VPS29A and VPS29B. Through a combination of AlphaFold predictive modelling, in vitro complex reconstitution, mass spectrometry and molecular cell biology, we find that the amino-terminal VPS29C extension constitutes an autoinhibitory sequence that limits access to a hydrophobic groove necessary for effector protein recruitment to Retromer, and association with Retriever and Commander. VPS29C is therefore unique in its ability to uncouple Retromer-dependent cargo sorting from the broader roles of VPS29A and VPS29B in regulating the endosomal-lysosomal network through accessory protein recruitment. Our identification and characterisation of VPS29C points to additional complexity in the differential subunit assembly of Retromer, an important consideration given the increasing interest in Retromer as a potential therapeutic target in neurodegenerative diseases.

**SIGNIFICANCE STATEMENT:** The endosomal-lysosomal network is essential for normal cellular function with network defects being associated with numerous neurodegenerative diseases. Two heterotrimeric complexes, Retromer and Retriever, control transmembrane protein recycling through the network. Of these, reduced Retromer expression is observed in Alzheimer’s disease and Retromer mutations lead to familial Parkinson’s disease. Here, we identify and characterise a new isoform of VPS29, a subunit shared between Retromer and Retriever. We reveal how this isoform, VPS29C, adopts an auto-inhibitory conformation to limit its association into Retriever and restrict the binding of VPS29C-containing Retromer to accessory proteins vital for regulating network function. By revealing added complexity in Retromer assembly and function, we provide new insight into Retromer’s potential as a therapeutic target in neurodegenerative diseases.

## INTRODUCTION

Across eukaryotes the establishment, maintenance, and adaptation of the functional cell surface proteome requires the selective sorting of internalised integral proteins through the intracellular endosomal network (Huotari and Helenius, 2011; Klumperman and Raposo, 2014; Cullen and Steinberg, 2018). The efficient sorting of channels, enzymes, transporters, receptors, adhesion molecules, and polarity cues is essential for an array of cellular, tissue and organism-level processes and physiology, with sorting defects being linked to developmental and age-related human diseases including cancer, metabolic syndromes and neurodegenerative diseases (Small et al., 2017; Sigismund et al., 2021; Gilleron and Zeigerer, 2023; Young et al., 2023).

Upon entering the endosomal network integral proteins and associated lipids and proteins (termed ‘cargo’) are sorted between two fates; either degradation within the lysosome or retrieval from this fate for recycling and reuse at the cell surface and other intracellular organelles (Cullen and Steinberg, 2018). While ESCRT complexes regulate the degradative fate (Migliano et al., 2022) a series of multi-protein assemblies orchestrate the retrieval and recycling pathways (McNally and Cullen, 2018; Chen et al., 2019). These include the Retromer, ESCPE-1, and WASH complexes (Seaman et al., 1998, Haft et al., 2000; Derivery et al., 2009; Gomez and Billadeau, 2009; Simonetti et al., 2019; Yong et al., 2020), the Retriever, CCC and Commander assemblies (Phillips-Krawczak et al., 2015; McNally et al., 2017; Healy et al., 2023; Boesch et al., 2024; Laulumaa et al., 2024), and additional sorting nexins (SNXs), SNX-BARs, and other proteins including ACAP1 (Carlton et al., 2004; Dai et al., 2004; van Kerkhof et al., 2005; Li et al., 2007; Strochlic et al., 2007; Traer et al., 2007; Lauffer et al., 2010; Harterink et al., 2011; Temkin et al., 2011; van Weering et al., 2012; Steinberg et al., 2013; Hsu et al., 2020).

Retromer and Retriever are stable heterotrimers assembled with a similar blueprint. For Retromer the amino-terminal end of the core VPS35 α-solenoid associates with VPS26A or VPS26B while its carboxy-terminal end binds to VPS29 (Hierro et al., 2007). Similarly, the amino-terminal end of the Retriever VPS35L α-solenoid binds to VPS26C, and the carboxy-terminal region binds to VPS29 (Healy et al., 2023; Boesch et al., 2024; Laulumaa et al., 2024). Retromer and Retriever, therefore, share VPS29 as a common subunit.

VPS29 adopts a phosphoesterase-fold and contains two conserved hydrophobic patches (Collins et al., 2005; Wang et al., 2005; Banos-Mateos et al., 2019). One associates with the carboxy-terminal regions of the VPS35 and VPS35L α-solenoids (Hierro et al., 2007; Healy et al., 2023; Boesch et al., 2024; Laulumaa et al., 2024), the other is solvent exposed and contains a surface groove. In Retromer, this groove is the primary site for binding to a variety of accessory proteins including TBC1D5, ANKRD27 and the FAM21 subunit of the WASH complex, that each present Pro-Leu motifs to occupy the groove (Gomez and Billadeau, 2009; Seaman et al., 2009; Harbour et al., 2010; Harbour et al., 2012; Jia et al., 2012; Hesketh et al., 2014; McGough et al., 2014a; McGough et al., 2014b Jia et al., 2016; Crawley-Snowdon et al., 2020; Guo et al., 2024; Romano-Moreno et al., 2024). In the case of Retriever, the VPS29 hydrophobic groove is occupied by an intramolecular Pro-Leu motif from the extended unstructured amino-terminal region of VPS35L (Healy et al., 2023; Boesch et al., 2024; Laulumaa et al., 2024). This intramolecular interaction stabilises the Retriever heterotrimer and its association with the CCC complex within the Commander super-complex and also acts to prevent Retriever from binding to Retromer accessory proteins (Healy et al., 2023; Butkovič et al., 2024). The function of the shared VPS29 subunit between Retromer and Retriever is therefore context dependent (Baños-Mateos et al., 2019).

Two human VPS29 isoforms have been identified that differ solely in their amino terminal sequences. The shorter form comprises ^1^MLVLVL^6^ (here termed VPS29A) while the longer form arises through inclusion of an additional short exon by alternative splicing and encodes four additional amino acids to give the sequence ^1^MAGHRLVLVL^10^ (here termed VPS29B). Both isoforms can assemble into Retromer but it has been suggested that the ^2^AGHR^5^ insertion modulates the affinity of VPS29 binding to the effector proteins TBC1D5 and AKNRD27 (Hesketh et al., 2014; Jia et al., 2016; Crawley-Snowdon et al., 2020). Here we identify a third human VPS29 isoform, VPS29C, with a significantly extended amino-terminal sequence of 28 amino acids. Using a combination of AlphaFold modelling, *in vitro* biochemical reconstitutions, and molecular cell biology approaches, we reveal that VPS29C preferentially assembles into the Retromer over the Retriever heterotrimer. We show that this stems from the 28-amino acid amino-terminal extension in VPS29C forming a transient intramolecular association that inhibits the hydrophobic groove necessary for VPS29 assembly into Retriever. In addition, this intramolecular association results in a VPS29C-Retromer complex that displays a limited ability to associate with Retromer accessory proteins that also depend on this VPS29 surface. Our identification and characterisation of VPS29C points to additional complexity in the differential subunit assembly of Retromer, an important consideration given the increasing interest in Retromer as a potential therapeutic target in neurodegenerative diseases (Small and Petsko, 2020).

## RESULTS

### Identification of a third VPS29 splice isoform

Two VPS29 isoforms, which we here term VPS29A and VPS29B, are ubiquitously expressed in human tissues and have been characterised at the molecular level (Haft et al., 2000; Collins et al., 2005; Damen et al., 2006; Jia et al., 2016; Crawley-Snowdon et al., 2020). These isoforms are generated by alternative splicing of the *VPS29* gene. We further noted a third computationally mapped alternative splicing variant of human *VPS29* (herein referred to as VPS29C, Gencode ID: ENST00000546588.1, UniProt ID: F8VXU5), which encodes an amino-terminal extension of 28 amino acids immediately prior to the ^2^AGHR^5^ insertion of VPS29B **(Figure 1A)**. An additional bioinformatic search revealed that highly similar extensions are found in a selection of primates, and more distantly related sequences are also present in several other mammalian lineages. This suggests that the VPS29C isoform likely evolved in the common ancestor of placental mammals after the divergence of placental mammals from marsupials **(Supplementary Table 1, Figure S1A-B)**.

**Figure 1.**
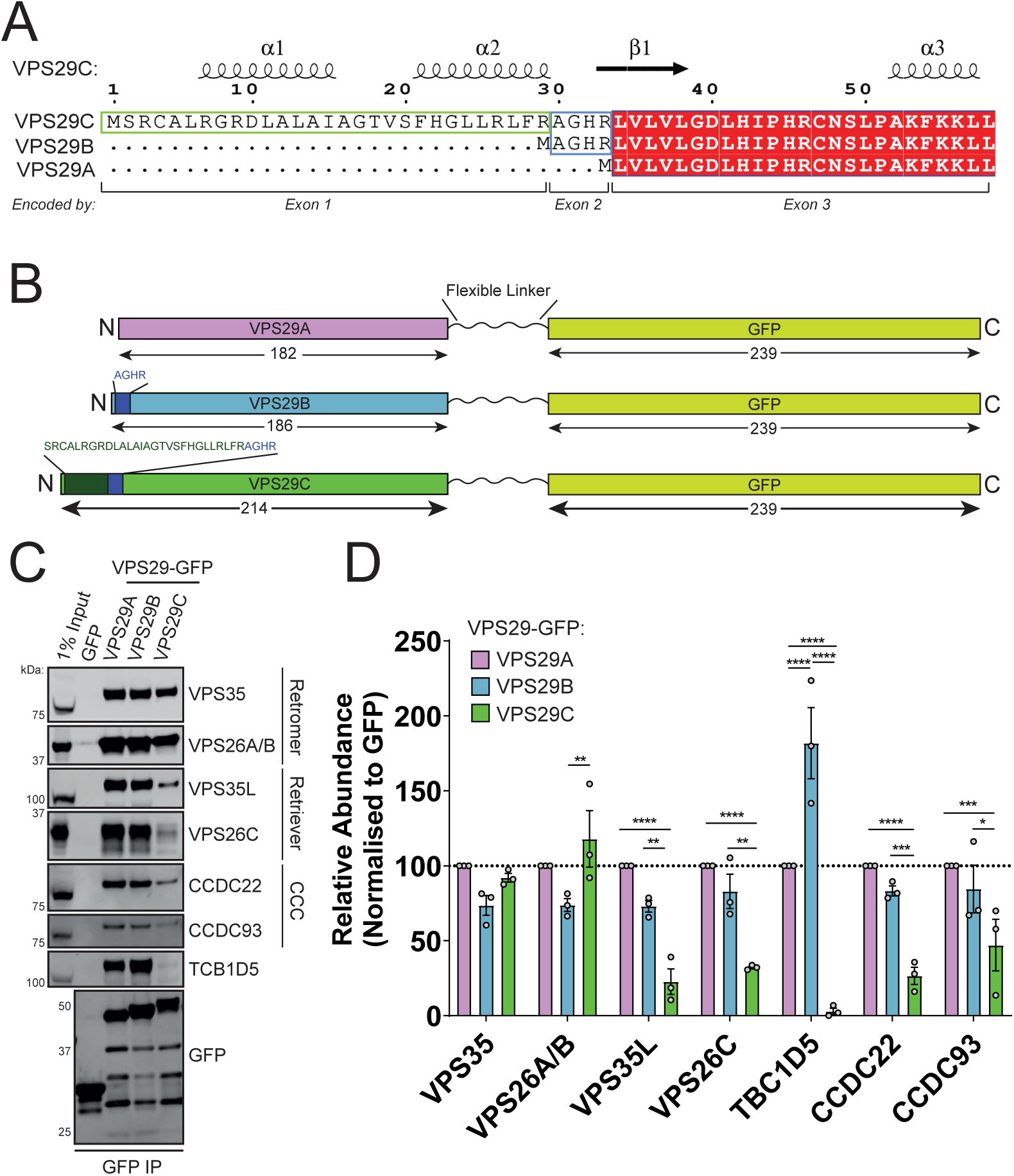
Identification of a novel VPS29 isoform that assembles into a functional Retromer complex. (A) Protein sequence alignment and corresponding exon junctions of human VPS29 splice isoforms reveal a unique amino-terminal extension loop in VPS29C. (B) Construct map describing the design of VPS29A/B/C-GFP constructs, with amino terminal extensions of VPS29B and VPS29C highlighted, and a flexible linker separating the carboxy-terminus of VPS29 from the amino-terminus of GFP. (C) GFP-nanotrap immunoprecipitation of VPS29A/B/C-GFP isoforms interaction with Retromer, Commander and TBC1D5. (D) Quantification of relative protein abundances in (C). Means +/- SEM, n = 3 independent experiments two-way ANOVA with Tukey’s multiple comparisons tests. * p < 0.05, ** p < 0.01, *** p < 0.001, **** p < 0.0001.

### VPS29C exhibits altered binding properties to VPS29A/B

To compare the biological functions of these VPS29 isoforms, VPS29A, VPS29B and VPS29C were carboxy-terminally tagged with green fluorescent protein (GFP) to leave their respective amino-terminal sequences unperturbed **(Figure 1B)**. Expression of VPS29A/B/C-GFP constructs in HEK293T cells followed by GFP-trap immunoprecipitation to isolate interacting proteins revealed a conserved ability of all three isoforms to associate with components of Retromer (VPS26A/B and VPS35) **(Figure 1C-D)**. VPS29A and VPS29B both robustly immunoprecipitated the well-documented VPS29 effector protein TCB1D5 with VPS29B showing a stronger association in agreement with *in vitro* biochemical data (Seaman et al., 2009; Hesketh et al., 2014; McGough et al., 2014; Jia et al., 2016; Crawley-Snowdon et al., 2020). Moreover, VPS29A and VPS29B also associated with components of the Retriever complex and Commander super-assembly with which VPS29 is known to interact (McNally et al., 2017; Healy et al., 2023; Boesch et al., 2024; Laulumaa et al., 2024). In comparison, VPS29C demonstrated significantly perturbed binding to the Retriever subunits VPS26C and VPS35L as well as components of the Commander complex CCDC22 and CCDC93. Moreover, VPS29C binding to TBC1D5 was also severely impaired **(Figure 1C-D)**.

### Interactome analysis is consistent with VPS29C displaying reduced association to accessory proteins and incorporation into Retriever

To expand our analysis, we performed an unbiased quantitative identification of the comparative VPS29A, VPS29B, and VPS29C interactomes. To establish this procedure, we first validated the expression and localisation of VPS29A/B/C constructs, each tagged with GFP at their carboxy-termini, by rescuing a previously validated VPS29 knockout (KO) H4 neuroglioma cell line using CRISPR-Cas9 guide RNAs that target all three VPS29 isoforms (Daly et al, 2023) **(Figure 2A).** Immunofluorescence analysis revealed that all three isoforms retained association with endosomes where they colocalised with endogenous VPS35 and the early endosomal marker EEA1 **(Figure 2B)**.

**Figure 2.**
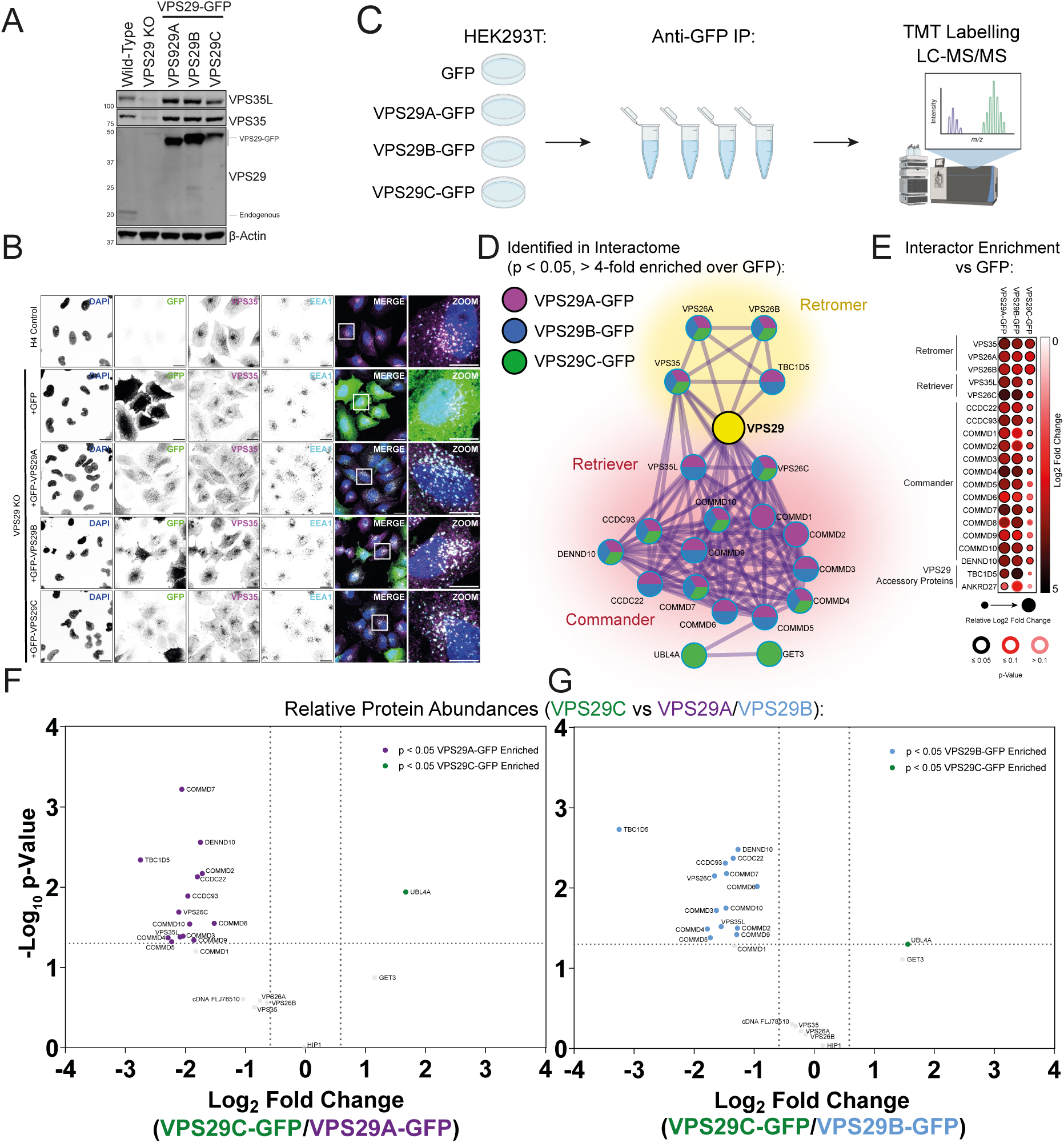
VPS29 isoforms exhibit different profiles of protein-protein interactions. (A) Stable H4 neuroglioma cell lines were made by rescuing VPS29 KO clonal cells with VPS29A/B/C-GFP. (B) Immunofluorescence microscopy of wild-type H4 cells and VPS29 KO H4 cells transfected with GFP, VPS29A-GFP, VPS29B-GFP or VPS29C-GFP. Cells were co-stained with anti-VPS35 and anti-EEA1 antibodies. Scale bar = 20µm, inset scale bar = 10µm. (C) Schematic of mass spectrometry experimental design. (D) Protein-protein interaction network of the isolated interactomes of VPS29A/B/C-GFP. Proteins are coloured by which VPS29-GFP isoform they were significantly identified by (Log_2_ fold change over GFP > 2, p < 0.05) as indicated in the legend. (E) Dotplots displaying relative enrichment of VPS29A/B/C-GFP interactors over GFP. Dot colour indicates Log_2_ fold change relative to GFP, dot size indicates relative fold change between VPS29 isoforms, and border colours indicate p value. (F-G) Volcano plots displaying the relative abundances of significant VPS29 interactors between isoforms following normalisation to GFP expression levels. Comparisons are made between VPS29C vs VPS29A (F) and VPS29C vs VPS29B (G) Significantly enriched or depleted proteins (Log_2_ fold change > 0.5, p < 0.05) are coloured.

We next expressed these constructs in HEK293T cells and performed GFP-trap immunoprecipitation followed by tandem mass spectrometry (TMT)-based proteomics to identify interactors of the three VPS29 isoforms **(Figure 2C)**. 19, 18 and 13 proteins were identified as significantly enriched by VPS29A-GFP, VPS29B-GFP and VPS29C-GFP, respectively (defined as proteins with > Log_2_ 2-fold enrichment over GFP-only condition, p < 0.05) **(Figure 2D, Figure S2A-C, Supplementary Tables 2-3)**. These interactors included almost all previously validated interactors of VPS29, including components of the Retromer, Retriever and Commander complexes, and TBC1D5 **(Figure 2D-E)**. In agreement with our biochemical data, VPS29C robustly interacted with the Retromer components VPS35, VPS26A and VPS26B, but demonstrated weaker associations with all other interactors including Retriever, the Commander complex, TBC1D5 **(Figure 2D-E)**. The association between VPS26C and ANKRD27 was also reduced, although ANRKD27 was only quantified in two experimental replicates (Figure 2E). In addition, UBL4A and GET3 were identified as uniquely enriched by VPS29C compared to VPS29A/B **(Figure 2D, Figure S2C)**. Notably, transmembrane Retromer and Retriever cargo proteins were not identified by any of the VPS29 isoforms, in line with previous findings of Retromer subunit immunoprecipitation combined with mass spectrometry (McMillan et al., 2016). This is likely due to the requirement for the coincidence detection of cargo, sequence-specific adaptors such as sorting nexin (SNX) proteins, and membrane lipid identity for Retromer and Retriever to robustly engage their transmembrane cargo.

To compare between VPS29 isoforms, the proteomics dataset was filtered for the proteins significantly interacting with at least one VPS29 isoform as defined in **Figure 2D**, then the abundances of interacting proteins were normalised to GFP, to control for differences in VPS29-GFP expression levels between conditions **(Supplementary Tables 4-5)**. The relative enrichments of proteins between VPS29 isoforms were then assessed, revealing that VPS29C-GFP displayed a significantly reduced association with almost all components of the Retriever and Commander complexes (VPS35L, VPS26C, CCDC22, CCDC93, COMMD1-10 and DENND10), and TBC1D5 when compared to VPS29A/B-GFP (< Log_2_ 0.5-fold change, p < 0.05) **(Figure 2F-G)** (Seaman et al., 2009; Hesketh et al., 2014; McGough et al., 2014; Jia et al., 2016; Crawley-Snowdon et al., 2020; Healy et al., 2023; Boesch et al., 2024; Laulumaa et al., 2024). Direct comparisons of significant interactors revealed minor differences between the interactomes of VPS29A-GFP and VPS29B-GFP, with VPS29B-GFP demonstrating a stronger enrichment of TBC1D5 in line with previous data and our quantitative Western blotting **(Figure 1C-D)**, although this did not reach statistical significance here **(Figure S2D)**. Taken together, these data define an interaction network for all three VPS29 isoforms and confirm that VPS29C demonstrates a reduced capacity to engage Retriever and associated Commander subunits, and most known Retromer interactors besides the core Retromer complex.

### Structural modelling of VPS29C reveals an extended helical amino terminus

To investigate the cause of the diminished VPS29C interaction network, we modelled the structure of VPS29C using AlphaFold2 implemented in ColabFold. The predicted local distance difference test (pLDDT) scores of 30 to 40 for the unique amino-terminal extension suggest that the region is predominantly unstructured **(Figure 3A)**. A similar prediction was also observed in models generated by AlphaFold3 and RoseTTAFold2 **(Figures S3A-B)**. Further analysis of the predicted aligned error (PAE) plot and multiple models generated from different seeds revealed that the unique amino-terminal extension is commonly predicted to form a short helical stretch positioned in proximity to the flat β-sheet surface of the VPS29 core **(Figures 3B-C, S3A-B)**. This hydrophobic groove is known to mediate binding to the Retromer accessory proteins TBC1D5, ANKRD27 and FAM21 (Hesketh et al., 2014; Jia et al., 2016; Crawley-Snowdon et al., 2020; Guo et al., 2024; Romano-Moreno et al., 2024) **(Figures 3F-G)**, as well as the bacterial protein RidL (Yao et al., 2018; Romano-Moreno et al., 2017; Barlocher et al., 2017). This same pocket is also essential for VPS29 incorporation into the Retriever complex, through binding to amino-terminal sequences in the VPS35L subunit (Healy et al., 2023; Boesch et al., 2024; Laulumaa et al., 2024). Alignment of this model with previously solved X-ray crystallographic structures of VPS29A and VPS29B revealed minimal deviation of the VPS29C model besides this amino-terminal extension **(Figures 3D-E)**.

**Figure 3.**
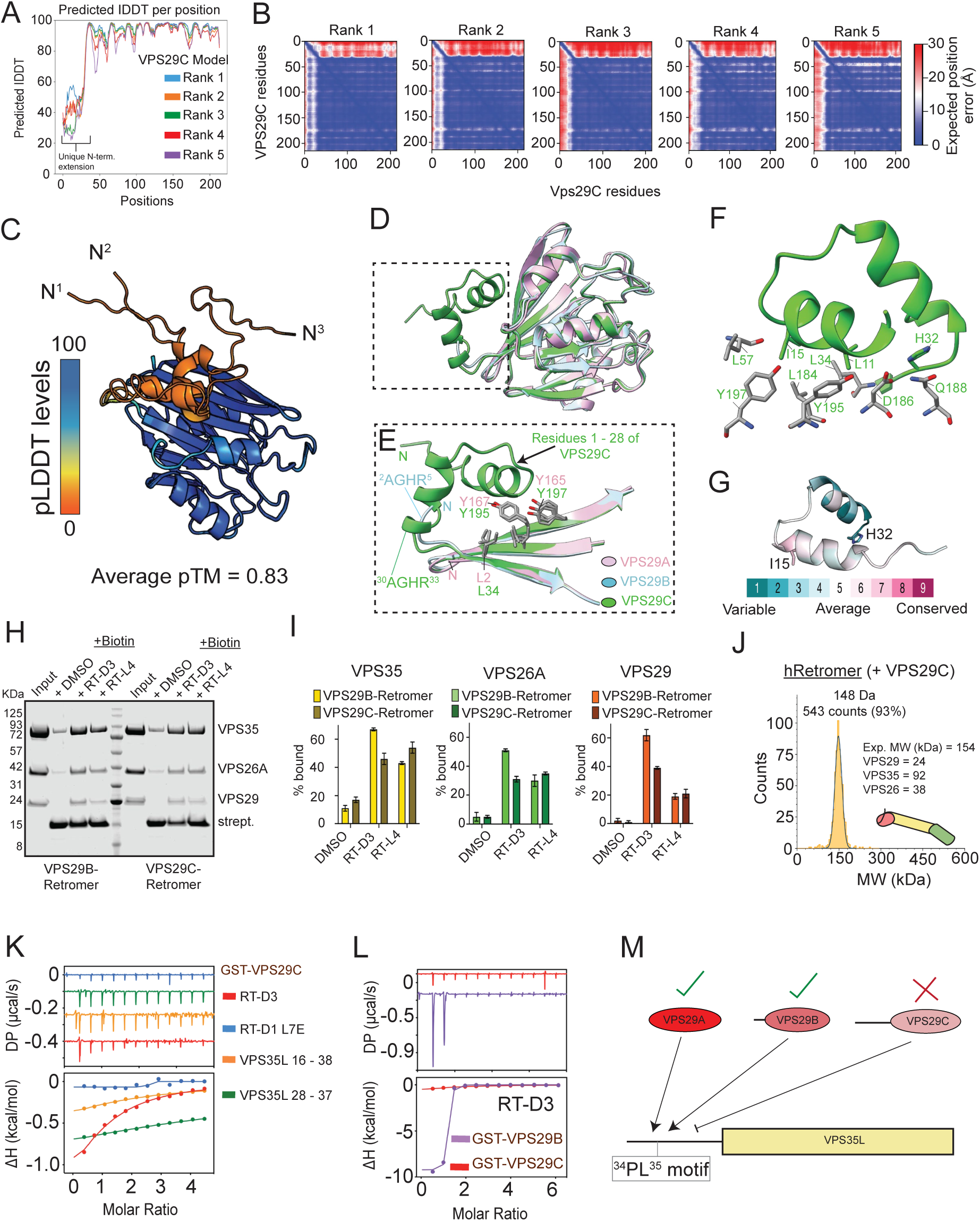
VPS29C displays an altered binding profile to other VPS29 isoforms. (A) AlphaFold2 pLDDT plot and (B) PAE plot of human VPS29C highlighting the flexibility of the extension loop. (C) Cartoon representation of AlphaFold2 predicted human VPS29C model. Three top ranked models are overlaid and coloured according to the pLDDT score. (D) Superposition of top ranked VPS29C model with the crystal structures of VPS29A (PDB ID: 5OSI) and VPS29B (PDB ID: 5GTU) to highlight the difference of the amino-terminal extension. (E) Highlight of the key binding region between the conserved hydrophobic pocket and the extension loop of VPS29C. (F) Highlight of the key interacting residues (in stick) of VPS29 shown in (E). (G) Sequence conservation map showing the conserved residues within the extension loop. (H) Pull-down assay of human Retromer containing VPS29B or VPS29C with streptavidin agarose coated with biotinylated RT-D3 or RT-L4. (I) Quantitation of the streptavidin-based pull-down assay shown in (H). Light yellow, green and red indicates the samples from VPS29B-containing Retromer. The heavy/dark yellow, green and red indicates the samples from VPS29C-containing Retromer. n = 2. (J) Mass photometry showing VPS29C-containing Retromer forming a 1:1:1 ratio complex. (K) ITC measurements of GST-VPS29C with RT-D3 (red), RT-D1 L7E (blue) and two different VPS35L Pro-Leu motif containing peptides (orange and green respectively). (L) ITC measurements of RT-D3 with either GST-VPS29B or GST-VPS29C to highlight the impact of the extension loop on binding to the Pro-Leu motif containing macrocyclic peptide known to show strong affinity to human VPS29A/B. All ITC graphs show the integrated and normalized data fit with a 1:1 ratio. (M) Schematic diagram summarising the binding capacity of each VPS29 isoform for VPS35L.

Looking more closely at the VPS29C model, we found the amino-terminal extension was typically oriented towards the VPS29 core due to the ^2^AGHR^5^ sequence forming a U-turn conformation as observed in the VPS29B structure **(Figure 3E)** (Jia et al., 2016). In three out of five models predicted by AlphaFold2, the hydrophobic residues within the amino-terminal extension including Leu11 and the conserved Ile15 reinforce this conformation through contact with Leu184 in the hydrophobic groove, which is equivalent to the Leu152/156 residue in VPS29A/B respectively that mediates accessory protein binding. We next compared the predicted structures of extended VPS29 isoforms from other mammalian species. While primate models appeared almost identical to the human VPS29C prediction in length and conformation (**Figure S4A)**, models from more distantly related mammals displayed variation in the length and structure of the amino-terminal extension yet were still regularly predicted to fold back over the hydrophobic groove in a similar way **(Figure S4B)**. These predictive models therefore suggest that the amino-terminal extension in VPS29C may form an intramolecular association with the potential to occlude access to the hydrophobic groove of VPS29 that is essential for binding to Retromer accessory proteins and in stabilising the Retriever heterotrimer.

### Recombinant VPS29C displays perturbed ligand binding

To validate that VPS29C could assemble into a Retromer heterotrimer, we first expressed and purified recombinant VPS35, VPS26A and VPS29B or VPS29C. Based on our previously published methods for purifying VPS29A- and VPS29B-containing Retromer complexes, we could similarly isolate the VPS29C-containing Retromer assembly (Chen et al., 2021). We next used a streptavidin-based pull-down using two biotin-tagged cyclic peptides known to associate with Retromer as the probe (Chen et al., 2021). For each cyclic peptide their incubation with the purified fractions revealed successful binding of all three proteins consistent with the formation of a stable VPS29C-containing Retromer heterotrimer complex (**Figures 3H** and **3I**). The nature of the VPS29C-Retromer complex was further examined by mass photometry which revealed a single peak with a molecular weight of 148 kDa, close to the expected molecular weight of Retromer (153.8 kDa) forming a 1:1:1 stoichiometric ratio complex (**Figure 3J**). Together, these independent biochemical approaches establish that VPS29C can efficiently assemble into a heterotrimeric Retromer complex like the VPS29A- and VPS29B-containing Retromer, supporting our immunoprecipitation data.

As discussed previously, VPS29A and VPS29B play crucial roles in the Retromer complex by dynamically recruiting effector proteins that modulate the dynamics of endosomal maturation such as TBC1D5, ANKRD27 and the WASH complex through FAM21 (Seaman et al., 2009; McGough et al., 2014; Jia et al., 2016; Crawley-Snowdon et al., 2020; Guo et al., 2024; Romano-Moreno et al., 2024), and in the Retriever complex by engaging the amino-terminus of the core VPS35L subunit (Healy et al., 2023; Boesch et al., 2024; Laulumaa et al., 2024). In these examples, binding occurs at a hydrophobic interface comprising a leucine residue (Leu-152 in VPS29A, Leu-156 in VPS29B), associated with a Pro-Leu motif in the interacting protein. In this mechanism, the ligand proline residue kinks the amino acid chain for optimal leucine engagement with the VPS29A/B binding pocket **(Figure S5)**. The Vps5 subunit of the *Saccharomyces cerevisiae* pentameric Retromer complex also engages Vps29 through this binding mechanism, indicating that this hydrophobic interface is evolutionarily conserved **(Figure S5)** (Chen et al., 2024; Shortill et al., 2024). Given that the corresponding hydrophobic interface in VPS29C appears to be at least partially occupied by the extended amino-terminus, we next investigated whether this may constitute an autoinhibitory mechanism that precludes ligand binding.

While we successfully isolated the VPS29C-Retromer using the streptavidin-based pull-down with high affinity cyclic peptides, we observed a subtle difference compared to the VPS29B-Retromer. Of the two cyclic peptides we used, RT-D3 presents a PL motif that specifically associates with high affinity to the Leu-containing hydrophobic groove in VPS29, such that it can outcompete binding to Retromer accessory proteins requiring this site, while the cyclic peptide RT-L4 binds to a region spanning the VPS26 interface with VPS35, without affecting the PL motif containing proteins from binding to VPS29 (Chen et al., 2021). RT-L4 was able to isolate both VPS29B-Retromer and VPS29C-Retromer equally, whereas RT-D3 was less efficient at isolating VPS29C-Retromer (**Figure 3H-I**). The reduced ability of RT-D3 to bind VPS29C may therefore be a consequence of the extended amino-terminus precluding access to Leu-184, while binding to other proteins such as VPS35 at distal interfaces is unaffected.

We further validated the ability of VPS29C to engage ligands by utilising isothermal titration calorimetry (ITC) to directly explore the binding affinity of cyclic peptides and natural peptides corresponding to the amino-terminal region of VPS35L, which contains a Pro-Leu motif that engages the VPS29 leucine pocket in the Retriever assembly (Healy et al., 2023; Boesch et al., 2024; Laulumaa et al., 2024). We observed binding of VPS29C to RT-D3 with a Kd of 13.4 μM while the RT-D1 L7E mutant control peptide, which we previously demonstrated has dramatically reduced affinity for VPS29, showed no binding as expected (**Figure 3K**). Although the PL motif-containing RT-D3 cyclic peptide can interact with VPS29C it has a significantly lower affinity compared to VPS29B (13.4 μM Kd for VPS29C vs 0.014 µM Kd for VPS29B) (**Figure 3K-L, Supplementary Table 6**). Consistent with these results and co-immunoprecipitation experiments we also found that VPS35L peptides harbouring a Pro-Leu motif showed only a weak binding affinity for VPS29C with a Kd >100 µM compared to the VPS29A data published previously (**Figure 3K, Supplementary Table 6**) (Healy et al., 2023). (**Figure 3L**). Overall, these experiments demonstrate the unique amino-terminal extension of VPS29C diminishes its ability to engage VPS35L when compared to VPS29A/B (**Figure 3M**).

### Mechanistic basis of VPS29C autoinhibition through an intramolecular interaction

Our examinations of VPS29C structural models identified hydrophobic residues, such as Leu-11 and Ile-15, that fold back towards Leu-184 in a conformation that resembles VPS29 ligand binding (**Figure 3D-G**, **Figure 4A**). The proximity of these hydrophobic residues may constitute an autoinhibitory mechanism that prevents the binding partners of VPS29 from accessing Leu-184, hence explaining the diminished binding of VPS29C to TBC1D5 and ANKRD27, and Retriever and Commander subunits (**Figure 2E**).

**Figure 4.**
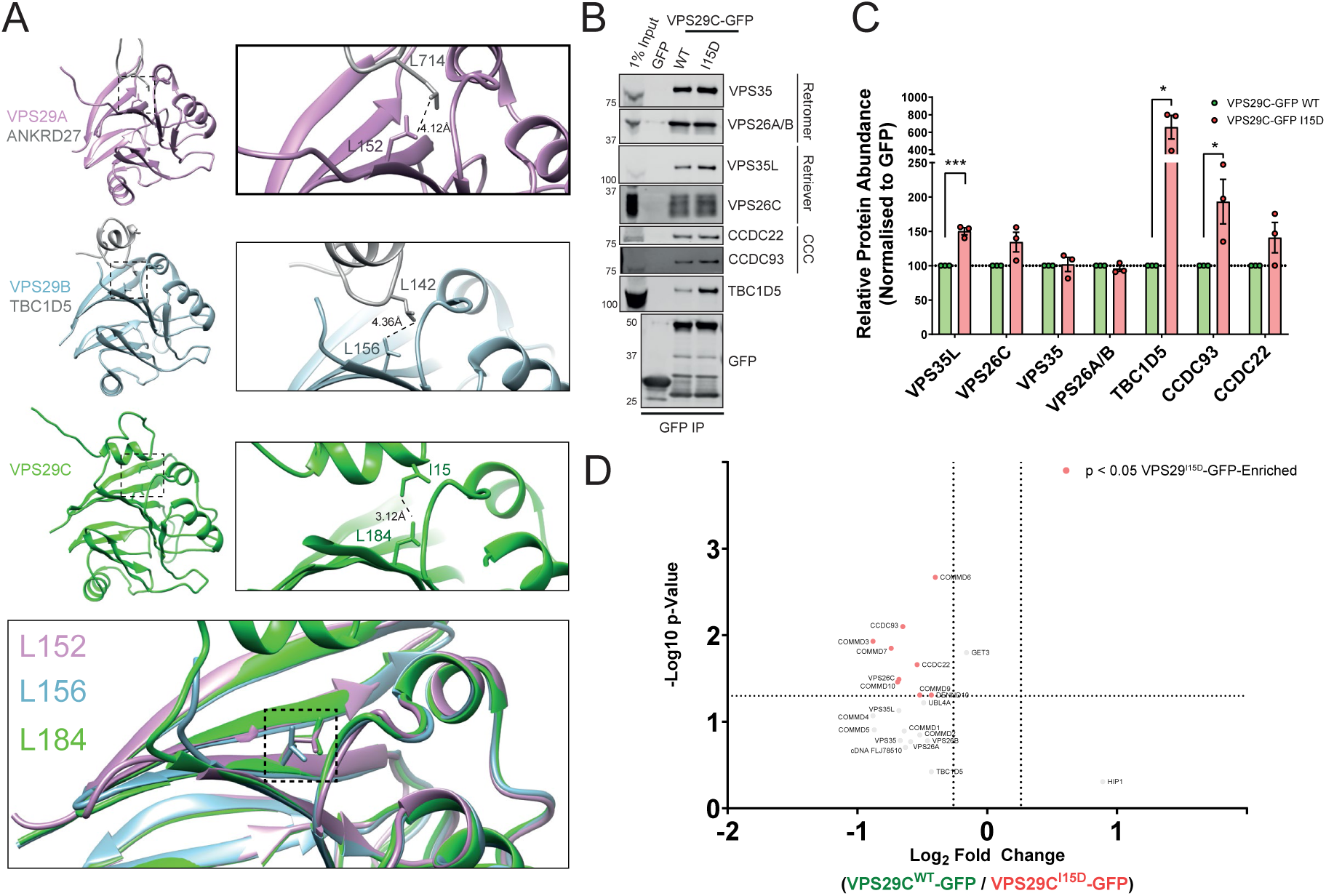
VPS29C is autoinhibited by an intramolecular interaction. (A) Comparison of VPS29A and VPS29B structures in complex with interacting regions of ANKRD27 (PDB: 6TL0) and TBC1D5 (PDB: 5GTU) respectively, with the VPS29C intramolecular interaction (AlphaFold model), and an overlay. The enlarged panels demonstrate the Leu152 (VPS29A) Leu156 (VPS29B) and Leu184 (VPS29C) residues involved in the binding interfaces. (B-C) GFP-nanotrap VPS29C^WT^-GFP and VPS29C^I15D^-GFP constructs expressed in HEK293T cells, followed by quantitative immunoblotting for VPS29 interactors. Means +/- SEM, n = 3 independent experiments, unpaired t-test. * p < 0.05, ** p < 0.01, *** p < 0.001, **** p < 0.0001. (D) Volcano plots displaying the relative abundances of significant VPS29 interactors between VPS29C^WT^-GFP and VPS29C^I15D^-GFP following normalisation to GFP expression levels, n = 3 independent experiments.

To test this hypothesis, we generated a VPS29C-GFP mutant with Ile-15 mutated to Asp (VPS29^I15D^) to introduce electrostatic charge into this hydrophobic sequence. Comparison of VPS29^WT^-GFP and VPS29^I15D^-GFP immunoprecipitations in HEK293T cells revealed an enhanced capacity for the mutant form to engage a selection of proteins, including significant enrichment of VPS35L, CCDC93 and TBC1D5, while binding to the core Retromer subunits VPS26A and VPS35 was unaffected **(Figure 4B-C)**. Furthermore, quantitative proteomic comparisons of VPS29^WT^-GFP and VPS29^I15D^-GFP interactors confirmed that this restoration of binding extends more widely to components of the Commander complex indicating a broader restoration of binding capacity, though TBC1D5 enrichment did not reach statistical significance through this approach **(Figure 4D)**. Together these data are consistent with the amino-terminal extension of VPS29C serving to block access to the hydrophobic Leu-184-containing interaction interface.

### VPS29C fails to regulate RAB7 dynamics and endosomal-lysosomal morphology

To explore the functional consequences of the altered associations of VPS29C, we utilised VPS29 KO H4 neuroglioma clone stably rescued with VPS29A/B/C-GFP constructs **(Figure 2A).** We recently characterised the widespread changes to endosomal-lysosomal morphology, identity and dynamics that occur because of deletion of the Retromer subunits VPS35 or VPS29 in H4 cells (Daly et al., 2023). We therefore investigated the ability of the VPS29-GFP constructs to reverse these phenotypes, beginning with cargo sorting. The glucose transporter GLUT1 is an established SNX27-Retromer cargo that undergoes endosome-to-plasma membrane trafficking through tubulovesicular carriers generated by the endosomal sorting complex for promoting exit-1 (ESCPE-1) (Steinberg et al., 2013; Simonetti et al., 2019; Yong et al., 2020; Yong et al., 2021; Simonetti et al., 2022). In VPS29 KO H4 cells, GLUT1 is strongly rerouted from the cell surface into LAMP1-positive compartments in the absence of sequence-dependent cargo sorting **(Figure 5A-B)**. Strikingly, this phenotype is rescued by re-expression of all three VPS29 isoforms, indicating that VPS29C assembles into a functional Retromer complex that can engage the SNX27 cargo adaptor and the ESCPE-1 complex to sort cargo away from a degradative fate.

**Figure 5.**
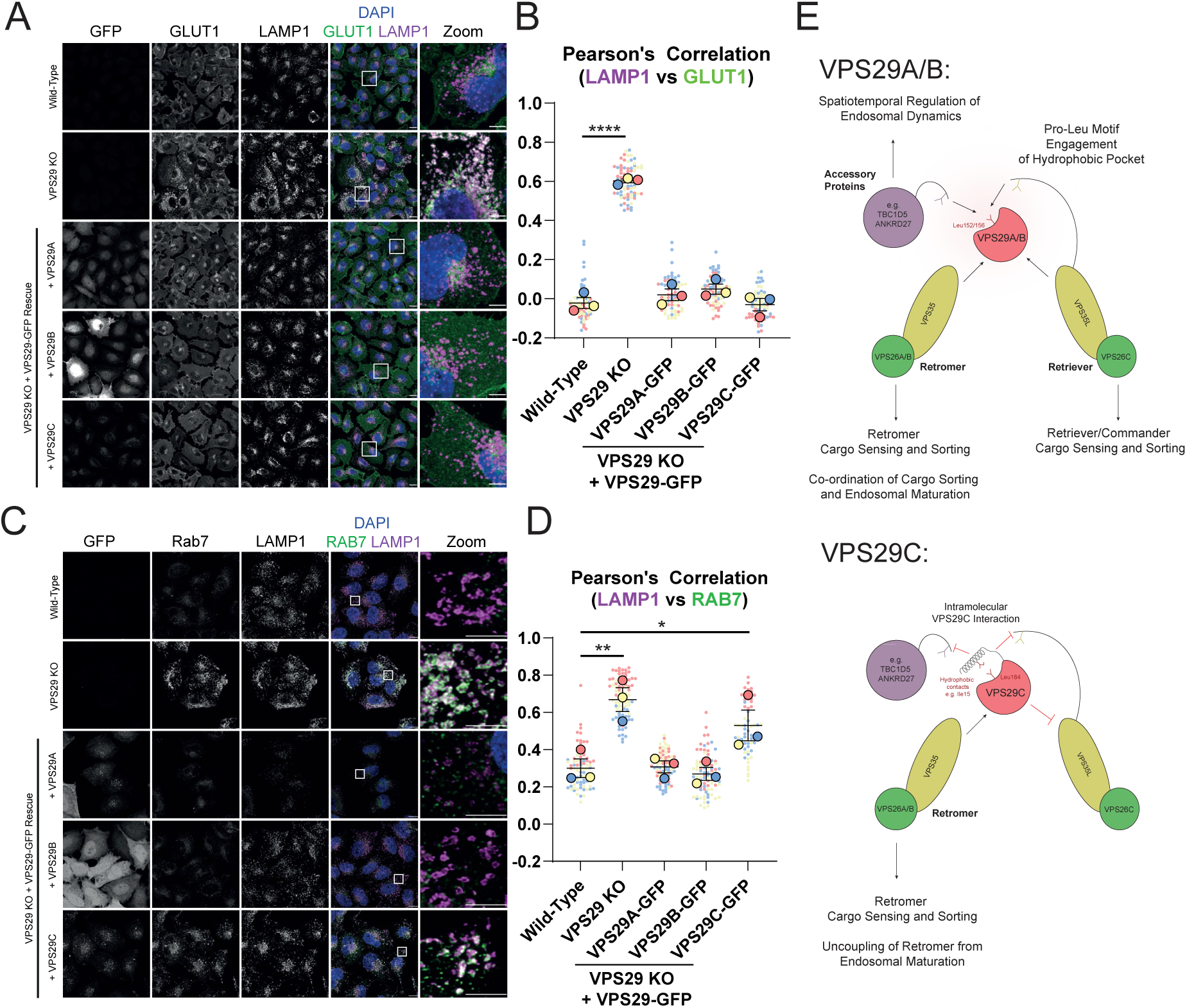
VPS29C-GFP can facilitate Retromer cargo sorting but is unable to regulate RAB7 Activity. (A) Immunofluorescence labelling of GLUT1 and LAMP1 in WT, VPS29 KO and VPS29A/B/C-GFP-expressing H4 cells. Scale bar = 20µm. (B) Quantification of Pearson’s correlation between GLUT1 and LAMP1. One-way ANOVA with Dunnett’s multiple comparisons tests, n = 3 independent experiments, * p < 0.05, ** p < 0.01, *** p < 0.001, **** p < 0.0001. (C) Immunofluorescence labelling of RAB7 and LAMP1 in WT, VPS29 KO and VPS29A/B/C-GFP-expressing H4 cells. Scale bar = 20µm. (D) Quantification of Pearson’s correlation between RAB7 and LAMP1. One-way ANOVA with Dunnett’s multiple comparisons tests, n = 3 independent experiments, * p < 0.05, ** p < 0.01, *** p < 0.001, **** p < 0.0001. (E) Model for the autoinhibitory function of the VPS29C amino-terminus in dictating Retromer and Retriever function.

It has previously been demonstrated that depletion of Retromer leads to RAB7 hyperactivation due to the lack of recruitment of the RAB7-GTPase-activating protein (GAP) and VPS29 binding partner TBC1D5 (Jimenez-Orgaz et al., 2018; Kvainickas et al., 2019). This is evidenced by the extensive recruitment of RAB7 to swollen LAMP1-positive compartments in VPS29 KO cells **(Figure 5C)**. As expected, re-expression of VPS29A/B-GFP restored this phenotype by limiting RAB7/LAMP1 colocalisation and resolving lysosomal morphology to a similar level to wild-type cells **(Figure 5C-D)** (Daly et al., 2022). In contrast, VPS29C-GFP, which binds TBC1D5 less efficiently, failed to rescue this phenotype despite its localisation on endosomal membranes **(Figure 5C-D)**. VPS29C is therefore able to uncouple the key functions of the Retromer complex as a master regulator of endosomal biology, by mediating cargo sorting but failing to control RAB7 GTP hydrolysis and effective endosomal-lysosomal maturation and resolution **(Figure 5E)**.

## DISCUSSION

Recent studies have illuminated the central role of VPS29 in orchestrating endosomal cargo sorting and dynamics through coordinating the Retromer, Retriever and Commander complexes (Jia et al., 2016, McNally et al., 2017, Jimenez-Orgaz et al., 2018, Healy et al., 2023; Boesch et al., 2024; Laulumaa et al., 2024). Here, we identify and characterise the biochemical properties of an additional isoform of VPS29, which we term VPS29C. This isoform contains an extended amino-terminal helical sequence that autoinhibits the hydrophobic groove on the VPS29 surface required for effector protein recruitment and assimilation into the Retriever complex. VPS29C therefore assembles preferentially into the Retromer complex to facilitate sequence-dependent cargo sorting but uncouples this activity from Retromer’s broader role in regulating RAB7 nucleotide cycling by its RAB7 GAP effector TBC1D5. Our recent work suggests that this subset of Retromer would therefore be less capable of regulating endosomal maturation, endosomal-lysosomal fusion and lysosomal reformation (Daly et al., 2023). Moreover, our proteomics data suggest that the extended amino-terminus of VPS29 does not appreciably expand its interaction network through gaining new interaction partners, but rather appears more likely to be autoinhibitory. Our biochemical data supporting this mechanism of autoinhibition, and its reversal through targeted site-directed mutagenesis, serve as an important functional validation of VPS29 activity and how its conducting role within the endosomal network is driven by accessibility of its interaction partners (**Figure 5E**).

Subunits of the Retromer and Retriever complexes are evolutionarily conserved amongst all eukaryotes. Notably, other ancient components of the endosomal sorting machinery have diversified into paralogues or isoforms that appear to play species- and tissue-specific roles. The modulation of endosomal sorting complexes through the selective incorporation of subunit isoforms or paralogues may provide context-dependent fine-tuning of endosomal-lysosomal dynamics. For example, alternative splicing of the *CLTC* gene to include an extra exon can modulate a switch between the formation of clathrin-coated pits to plaques in a tissue-specific context (Guidice et al., 2014; Moulay et al., 2020). Additionally, the ESCPE-1 subunit SNX32 is a paralogue of SNX6 that is predominantly expressed in the brain where it may engage neuronal cargoes (Sugatha et al., 2023). Three paralogues of VPS26 are expressed in humans, termed VPS26A, VPS26B and VPS26C, of which the former two assimilate into Retromer and the latter into Retriever (Kerr et al., 2005, Collins et al., 2008, McNally et al., 2017). The VPS26B gene emerged in chordates through duplication of VPS26A, whereas VPS26A and VPS26C are conserved amongst eukaryotes (Burgacic et al., 2011). VPS26A and VPS26B predominantly differ in their carboxy-terminal sequences, and while they exhibit almost identical interaction networks in human retinal pigment epithelial cells (McMillan et al., 2016), differential cargo engagement by these proteins has been suggested (Kim et al., 2010, Bugarcic et al., 2011, Bugarcic et al., 2015). Mutations associated with atypical parkinsonism are found in VPS26A specifically (Gustavsson et al., 2015, McMillan et al., 2016) and conversely VPS26B-Retromer is specifically enriched in the *trans*-entorhinal cortex region of the brain that is particularly vulnerable to Alzheimer’s disease-related neurodegeneration (Simoes et al., 2021), emphasising that spatially coordinated expression of subunit-specific endosomal sorting complexes and engagement of tissue-specific cargoes may be crucial to neuroprotection.

In contrast to the widespread conservation of the Retromer and Retriever complexes, the extended VPS29C amino-terminal sequences appear to have arisen relatively recently in evolutionary history and have apparently been independently lost on a number of occasions. AlphaFold modelling suggests that these mammalian amino-terminal extensions are, like the human sequence, also predicted to fold in over the conserved VPS29 hydrophobic groove in a conformation that resembles the autoinhibited state of VPS29C. These models raise the interesting possibility that the modulation of VPS29 activity through alternative splicing arose following the divergence of placental mammals from marsupials, though substantiating this hypothesis requires future structural and biochemical validation. The tissue expression profiles of VPS29A, VPS29B and VPS29C remain to be clarified, but it is conceivable that similar subtle differences of VPS29 isoform localisation and cargo engagement may occur across tissues. Spatially mapping VPS29C expression therefore represents an avenue of future work that may illuminate its functional context.

In conclusion, this study characterises an additional VPS29 isoform that regulates effector binding through autoinhibition. The biochemical inhibitory mechanism we elucidate serves to validate the central orchestrating role of VPS29 in the endosomal network. Given that the concerted activity of VPS29-associated endosomal sorting complexes are broadly considered neuroprotective, approaches to modulate their activity represent promising future therapeutic avenues. The autoinhibitory VPS29C isoform we describe here therefore adds further evidence to the notion that activity of these crucial proteins can be fine-tuned and may inform the future development of therapeutic strategies to re-balance Retromer activity.

## Supporting information

Proteomics Guide

## FIGURES AND FIGURE LEGENDS

**Figure S1.**
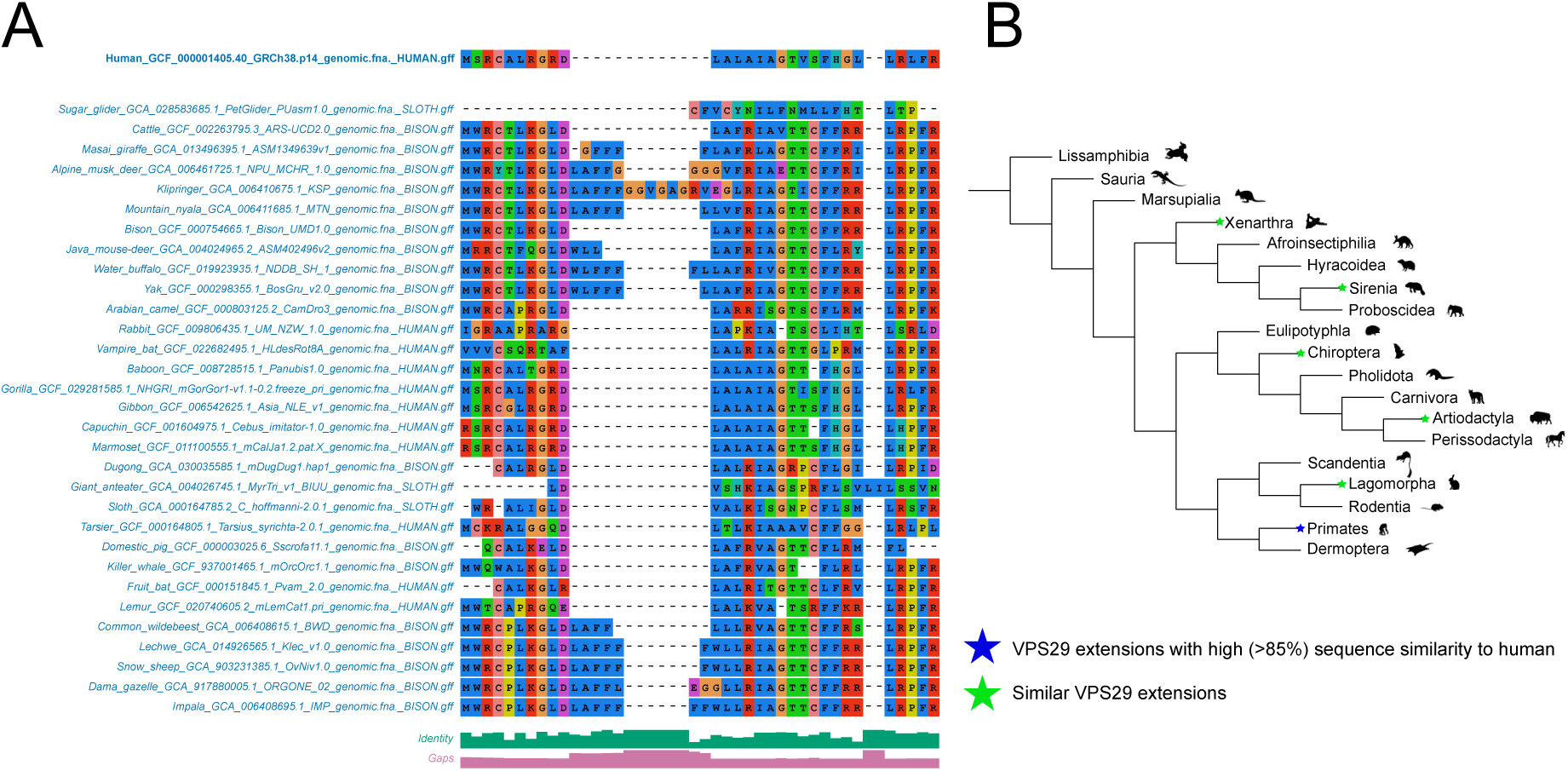
Phylogeny of organisms with an extended VPS29 amino terminus. (A) Sequence alignment of animal VPS29 extended isoforms. The identifier for each sequence contains the common name of the species encoding the isoform, the genome accession number, and the query sequence (from human, bison or sloth) used to identify the isoform. (B) Schematic phylogenetic tree of animal evolution. Orders containing species with identified VPS29 extended amino-termini are labelled green or blue depending on their degree of sequence similarity to the human VPS29C sequence.

**Figure S2.**
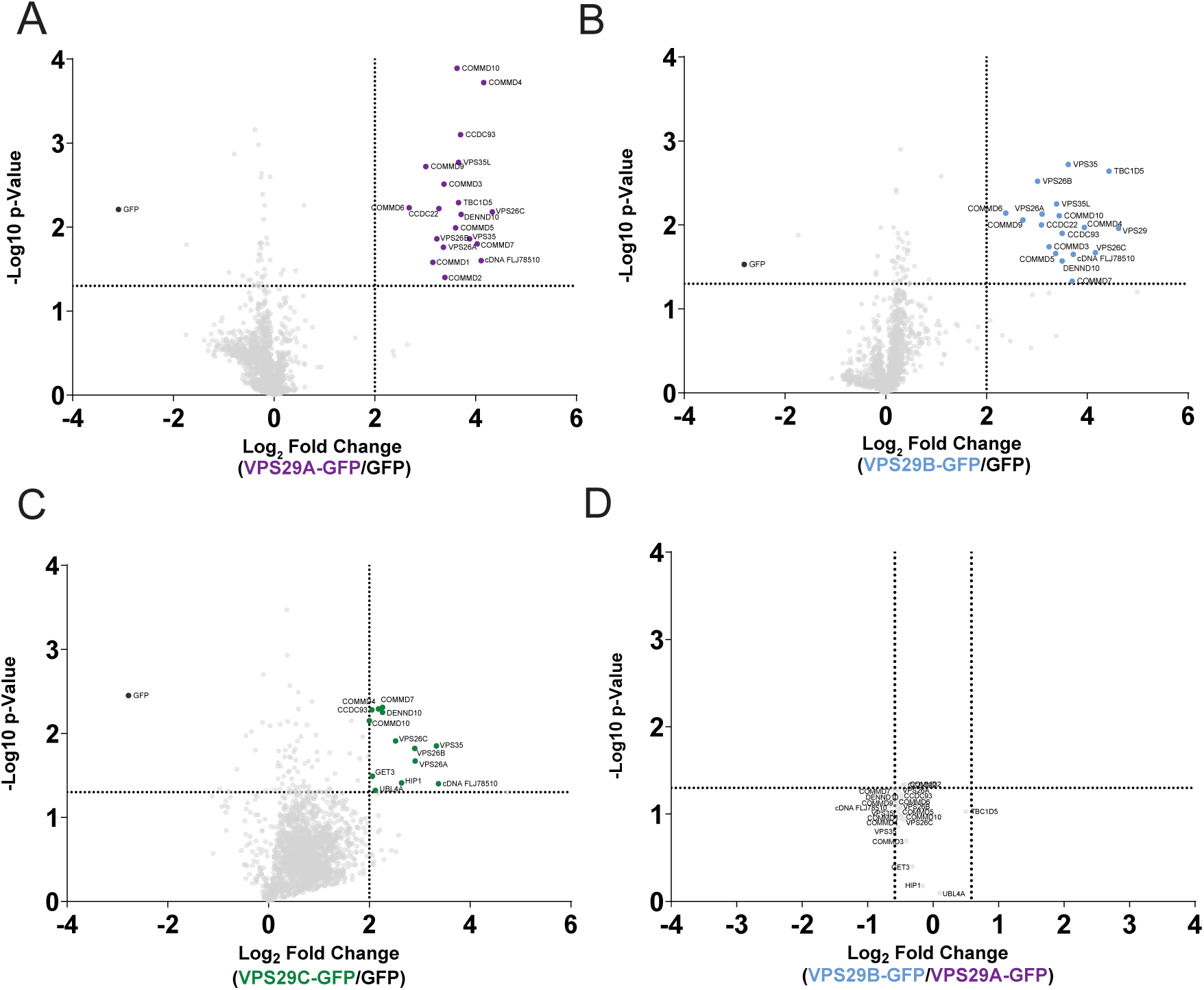
Identification of VPS29 interactors through quantitative proteomics. (A-C) Volcano plots displaying relative enrichment of proteins by VPS29A (A), VPS29B and VPS29C (C) over GFP. Proteins were defined as significant if enrichment exceeded Log_2_ fold change > 2, and p < 0.05. n = 3 independent experiments. (D) Volcano plot displaying the relative abundances of significant VPS29 interactors between VPS29A and VPS29B following normalisation to GFP expression levels.

**Figure S3:**
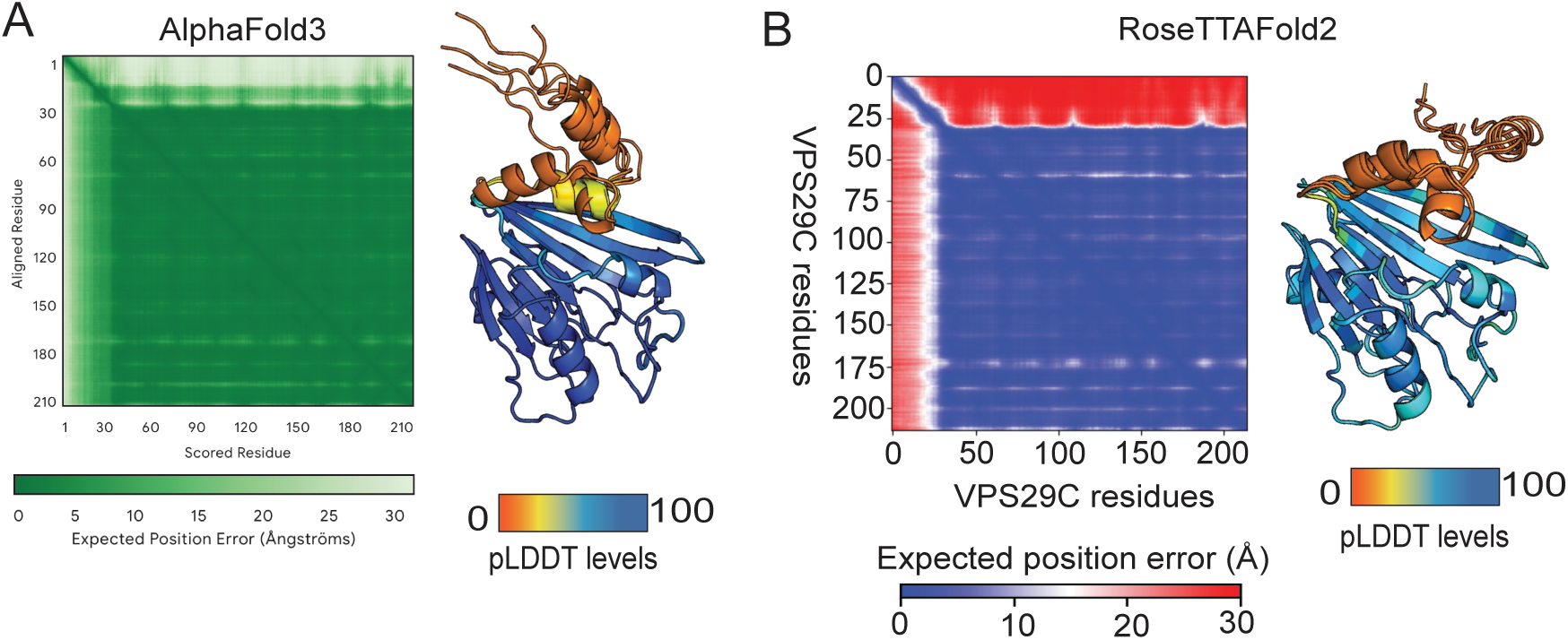
**Structural models of VPS29C generated by AlphaFold3 and RoseTTAFold2**. (A) PAE plot and cartoon representation of AlphaFold3 and (B) RoseTTAFold2 predicted human VPS29C model. In both cases, top ranked models are overlaid and coloured according to the pLDDT score

**Figure S4.**
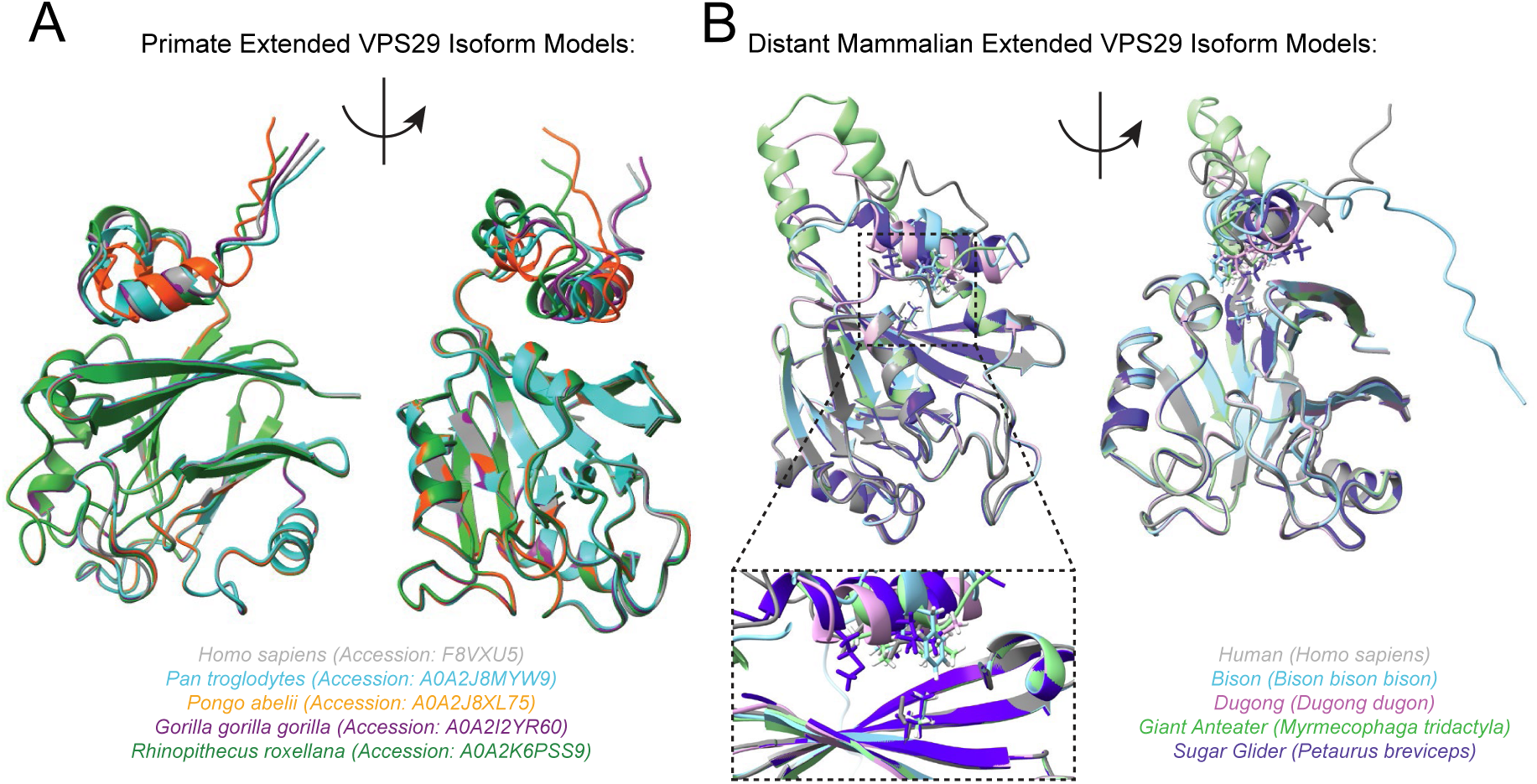
Structural prediction of related VPS29 sequences with extended amino termini. (A) Overlay of AlphaFold2 models of primate VPS29C isoforms, with their corresponding Uniprot accession IDs labelled. (B) Overlay of AlphaFold2 models of human and mammalian VPS29 sequences with extended amino termini. Hydrophobic residues in proximity of Leu-184 or the equivalent Leu residue are shown in stick formation.

**Figure S5.**
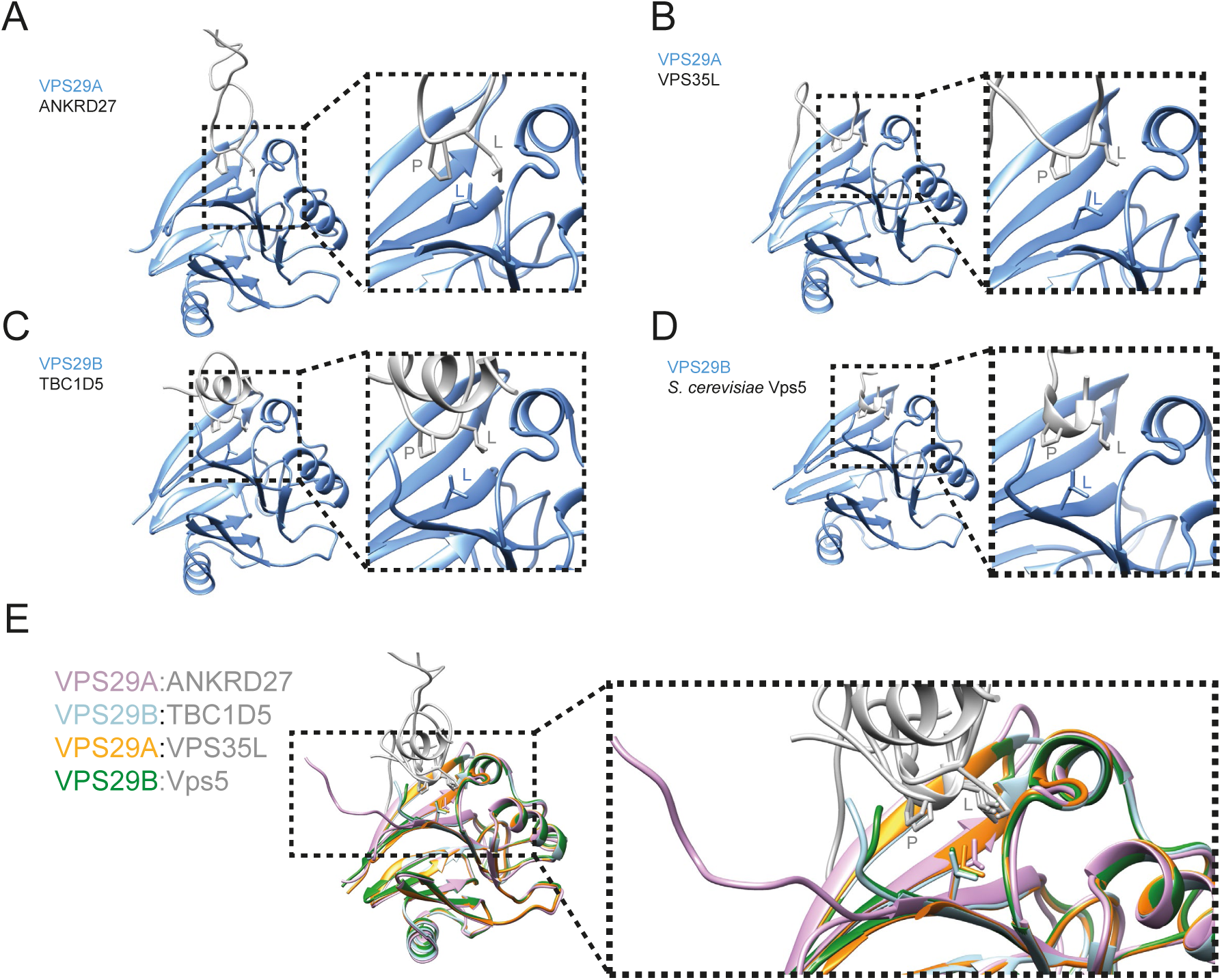
Evolutionary conservation of VPS29 binding to Pro-Leu motif ligands. (A-D) Protein structures of VPS29 isoforms bound to the interacting proteins VARP (PDB: 6TL0), TBC1D5 (PDB: 5GTU), VPS35L (PDB: 8SYL and 8ESE) and *Saccharomyces cerevisiae* Vps5 (PDB: 8FUD). (E) Overlay of structures displayed in (A-D) demonstrating structural similarity of the Pro-Leu motif binding mechanism.

## Supplementary Tables

**Supplementary Table 1**

Protein BLAST output of the VPS29C amino terminal extension amino acid sequence. Similar sequences in other organisms are ranked by identity matching.

**Supplementary Table 2**

Raw tandem mass tagging (TMT) proteomics data of GFP, VPS29A-GFP, VPS29B-GFP, VPS29C-GFP and VPS29C^I15D^-GFP immunoprecipitates. Protein log_2_ fold changes are presented as relative to their abundance in the GFP only control condition. *N* = 3 independent experiments.

**Supplementary Table 3**

Lists of proteins from Supplementary Table 2 that were significantly enriched (Log2 fold change > 2 relative to GFP, > 1 unique peptide, p < 0.05) by either VPS29A-GFP, VPS29B-GFP or VPS29C-GFP.

**Supplementary Table 4**

TMT proteomics data of GFP, VPS29A-GFP, VPS29B-GFP, VPS29C-GFP and VPS29C^I15D^-GFP immunoprecipitates normalised to GFP levels. Protein log_2_ fold changes are presented as relative abundances between VPS29-GFP isoform constructs. *N* = 3 independent experiments.

**Supplementary Table 5**

Lists of proteins from Supplementary Table 4 that were significantly enriched or depleted (Log2 fold change +/- 0.5 relative to different VPS29 isoforms, > 1 unique peptide, p < 0.05) by either VPS29A-GFP, VPS29B-GFP or VPS29C-GFP.

**Supplementary Table 6**

Thermodynamic parameters for the binding of Vps29C, Vps35L and cyclic peptides by ITC.

## ACKNOWLEDGEMENTS

We thank the Wolfson Bioimaging Facility at the University of Bristol for their support. J.L.D. is supported by a Wellcome Early Career Award (225128/Z/22/Z). R.B. is supported by the EndoConnect European Research Training Network (No. 953489). Work in the Cullen laboratory is supported by the Wellcome Trust (104568/Z/14/Z and 220260/Z/20/Z), the Medical Research Council (MR/L007363/1 and MR/P018807/1), the Lister Institute of Preventive Medicine, and the award of a Royal Society Noreen Murray Research Professorship to P.J.C. (RSRP/R1/211004). B.M.C. is supported by an Investigator Grant, Senior Research Fellowship and Project Grant from the National Health and MRC (APP2016410, APP1136021 and APP1156493).

## AUTHOR CONTRIBUTIONS

Biochemistry and cell biology analysis: J.L.D., K.J.M., R.B., E.P., G.G-S., Z.H. and P.J.C. Biophysics and AlphaFold2: K-E.C., Q.G. J.L.D and B.M.C. Proteomics and Bioinformatic analysis: J.L.D, P.A.L. and K.J.H. Evolutionary analysis: E.R.R.M. and T.A.W. Manuscript Writing - 1^st^ draft: J.L.D., K.J.M. B.M.C. and P.J.C; Final Version: all authors. Initial Concept: J.L.D., K.J.M., B.M.C. and P.J.C. Concept Development: all authors. Funding and Supervision: J.L.D., K.J.M., T.A.W., B.M.C. and P.J.C.

## METHODS

### Materials and Methods

#### Antibodies

Primary antibodies include: β-Actin (Sigma-Aldrich; A1978; clone AC-15; 1:2000 Western blot (WB)), CCDC22 (Proteintech, 16636-1-AP, 1:1000 WB), CCDC93 (LS Bio, C336997, 1:1000 WB), EEA1 (Cell Signalling, 3288S, 1:200 immunofluorescence (IF)), GFP (Roche; 11814460001; clones 7.1/13.1; 1:1000 WB; 1:400 IF), GLUT1 (Abcam; EPR3915; ab115730; 1:200 IF), LAMP1 (Developmental Studies Hybridoma Bank; AB_2296838; clone H4A3; 1:400 IF) LAMP1 (Abcam; ab21470; 1:200 IF), RAB7 (Abcam; EPR7589; ab137029; 1:200 IF), TBC1D5 (Abcam, 204896, 1:1000 WB), VPS26A (Abcam, ab137447, 1:1000 WB), VPS26C (Merck, ABN87, 1:1000 WB), VPS29 (Santa Cruz; D-1; sc-398874; 1:500 WB), VPS35 (Abcam; ab157220; clone EPR11501(B); 1:1000 WB), VPS35 (Abcam, ab10099, 1:200 IF), VPS35L (Abcam, ab97889, 1:1000 WB)

Secondary antibodies: For Western blotting, 680nm and 800nm donkey anti-mouse and anti-rabbit fluorescent secondary antibodies (Invitrogen, A-21057, A3275 - 1:20,000). For immunofluorescence, 488nm, 568nm and 647nm AlexaFluor-labelled anti-mouse,anti-rabbit and anti-goat secondary antibodies (Invitrogen, A32753, A32731, A10037, A10042, A32787, A-11057-1:400). 0.5 µg/mL 4’, 6-diamidino-2-phenylindole dihydrochloride (DAPI; Sigma-Aldrich, D8417) was added to secondary antibody mixtures to label DNA.

#### Cell Culture

HEK293T cells were sourced from the American Type Culture Collection (ATCC). H4 neuroglioma cells were a gift from Dr Helen Scott and Professor James Uney (University of Bristol). Clonal VPS29 KO cells were generated by transfecting cells with Cas9 and guide RNA sequences targeting the sequences 5’-GGACATCAAGTTATTCCAT-3’ and 5’-GGCAAACTGTTGCACCGGTG-3’ within *VPS29* and validated previously (Daly et al., 2023).

Cells were grown in Dulbecco’s Modified Eagle Medium (DMEM; Sigma-Aldrich), supplemented with 10% (vol/vol) fetal bovine serum (FBS) (Sigma-Aldrich) and penicillin/streptomycin (Gibco). H4 cells were transduced with human immunodeficiency virus (HIV)-1-based lentiviruses for stable expression (construct of interest in pLVX-GFP-IRES-Puro plasmid backbone, and pCMV-dR8.91 packing plasmid) pseudotyped with vesicular stomatitis virus (VSV)-G envelope plasmid (pMDG2). HEK293T cells were transfected with the constituent plasmids using polyethyleneimine (PEI) transfection, then lentiviruses were harvested after 48 hours. H4 cells were seeded into a plate, then transduced with lentivirus following adherence. 3 μg/mL puromycin dihydrochloride was used for selection of VPS29-GFP-expressing cells.

#### Synthesis and Cloning of VPS29 isoforms

The protein-coding sequences of VPS29A, VPS29B and VPS29C followed by a flexible linker encoding the sequence GGGGSGGGGS were synthesised as dsDNA fragments (Eurofins Genomics) with flanking restriction to produce the cassette (*EcoRI*)-VPS29-(*KpnI*)-Linker-(*BamHI*), then digested with EcoRI and BamHI and ligated into GFP-containing the GFP-containing backbones pEGFP-N1 for transient transfection, or pLVX-GFP-IRES-Puro for stable lentiviral transduction.

#### Quantitative Western Blotting

NuPAGE 4-12% gradient Bis-Tris precast gels (Life Technologies, NPO336) were used for SDS-PAGE, followed by transfer onto methanol-activated polyvinylidine fluoride (PVDF) membrane (Immobilon-FL membrane, pore size 0.45 μm; Millipore, IPFL00010). Membrane was blocked, then sequentially labelled with primary and secondary antibodies. Fluorescence detected by scanning with a LI-COR CLS Odyssey scanner and Image Studio analysis software (LI-COR Biosciences) was used to quantify band intensities.

#### Immunofluorescence Microscopy and Analysis

H4 cells were transfected with GFP or VPS29-GFP constructs using Fugene 6 (Promega) or stably expressed using lentiviral vectors. H4 cells were seeded onto 13 mm coverslips the day before fixation. DMEM was removed, followed by two washes with PBS, then cells were fixed in 4% paraformaldehyde (PFA) (Pierce, 28906) for 20 minutes at room temperature. Cells were permeabilised in either 0.1% Triton X-100 or 0.1% (w/v) saponin (Sigma-Aldrich, 47036) for 5 minutes followed by blocking with 1% (w/v) BSA, (plus 0.01% saponin for those permeabilised by saponin) in PBS for 15 minutes. Coverslips were stained with primary antibodies for 1 hour, followed by secondary antibodies for 30-60 minutes, then mounted onto glass microscope slides with Fluoromount-G (Invitrogen, 00-4958-02).

Confocal microscope images were taken on either a Leica SP8 confocal laser scanning microscope attached to a Leica DM l8 inverted epifluorescence microscope or a Leica SP5-II Biosystems confocal laser-scanning microscope attached to a Leica DMI6000 inverted epifluorescence microscope (Leica Microsystems), with a 63x UV oil immersion lens, numerical aperture 1.4 (Leica Microsystems, 506192), and acquired using LAS AF software (Leica Microsystems). Colocalisation and fluorescence intensity analysis was performed using Volocity 6.3 software (PerkinElmer) with automatic Costes background thresholding. Immunofluorescence images were prepared in either Image J (Fiji) or Volocity 6.3.

#### GFP Immunoprecipitation

HEK293T cells were transfected with GFP or VPS29-GFP constructs using polyethylenimine (PEI), then lysed 48 hours later with lysis buffer (50mM Tris-HCl, 0.5% NP-40 PBS with 1 x protease inhibitor cocktail (Roche)). Lysates were spun at 18,000 x g for 10 minutes at 4°C, then the supernatants were transferred onto GFP-trap beads (Proteintech, GTA20) and incubated for 1 hour at 4°C. Beads were then washed twice with 50mM Tris-HCl, 0.25% NP-40 PBS wash buffer, then once with wash buffer without NP-40 prior to elution in 2X LDS sample buffer (Invitrogen), 1.5% β-mercaptoethanol.

#### Recombinant protein expression and purification in bacteria

For the expression of human Retromer in bacteria, full-length cDNA encoding human VPS35, human VPS26A, human VPS29C and mouse VPS29B were cloned into either N-terminal His-tagged pET28a or GST-tagged pGEX6P1 vectors.

All the bacterial constructs were expressed in BL21 (DE3) competent cells using the standard autoinduction method as described previously. Cell cultures were harvested by centrifugation at 6000 X g for 5 min at 4°C and the cell pellets were then stored at

-80°C until cell lysis. To obtain individual Retromer subunit including VPS26A, GST-VPS29B, and GST-VPS29C, the frozen cell pellet was resuspended in protein buffer (50 mM Tris-HCl pH 7.5, 200 mM NaCl, 2 mM β-mercaptoethanol) supplemented with 50 μg/mL benzamidine, 100 units of DNase I. In all cases, the resuspended cells were lysed through a Constant System TS-series high pressure cell disruptor. The soluble homogenate cleared by centrifugation was loaded onto either Talon resin (Clontech) for His-tagged VPS26A or Glutathione Separose (GE Healthcare) for GST-tagged VPS29B/VPS29C. The fusion-tagged containing proteins eluted from the resin were then further purified using exclusion chromatography (SEC) on a HiLoad® 16/600 Superdex 200 or Superdex 75 column equilibrated with protein buffer. For Retromer, the soluble homogenate was first loaded onto Talon resin (Clontech) followed by glutathione Sepharose (GE healthcare) to obtain the correct stoichiometry ratio of the Retromer complex. Removal of the GST tag from the constructs were performed using on-column cleavage approach with PreScission protease. The fusion tag free protein was further purified using SEC on a HiLoad® 16/600 Superdex 200 column as described above. Protein integrity was validated using SDS-PAGE gel and concentration was measured using Nanodrop at OD280.

#### Isothermal Titration Calorimetry

All microcalorimetry experiments were carried out at 25°C using a PEAQ ITC (Malvern, UK) in protein buffer. The interaction of VPS29 and peptides were carried out by titrating 300 μM of RT-D3, RT-D1 L7E, VPS35L 28 - 37 or VPS35L 16 – 38 into 10 μM of GST-tagged VPS29C or GST-tagged VPS29B. In the macrocyclic peptide experiment, the equivalent percentage (v/v) of DMSO was added into the target protein to avoid buffer mismatch. All ITC experiments were performed with a single injection of 0.4 μL followed by a series of 12 injections of 3.2 μL each, spaced 180 seconds apart, and stirred at 750 rpm. To calculate the heat exchange of interactions, the observed peaks were integrated with the subtraction of heat of dilution from the background. The thermodynamic parameters including Kd, ΔH, ΔG, and −TΔS were obtained by fitting and normalized the data to a single-site binding model using Malvern software package. The stoichiometry was adjusted initially, and if the value was near 1, N was set to exactly 1.0 for computation. To ensure the data were reproducible, each experiment was run at least twice.

#### Mass photometry

Molecular mass measurement of VPS29C containing Retromer was performed using a Refeyn OneMP mass photometer (Refeyn Ltd) following manufacturer’s instructions. For the measurement to take place, the purified Retromer in protein buffer at a final concentration of 50 nM - 100 nM was loaded onto the mass photometer, and 1,000 frames were recorded. The data was examined using Refeyn DiscoverMP software. The molecular mass was calculated using the calibration curves of protein standards with known molecular weight (ie. 66, 132 and 440 kDa) in the same protein buffer.

#### Biotinylated cyclic peptide pull-down assay

Streptavidin-based pull-down assay was carried out by adding the purified Retromer fraction with either biotinylated RT-D3 or biotinylated RT-L4 prior of loading into the streptavidin agarose. As a negative control, equal amount of purified Retromer fraction was also loaded directly into the streptavidin agarose without macrocyclic peptide. To avoid precipitation caused by the macrocyclic peptides, the Retromer – peptides mixtures were centrifuged at 17,000 rom for 20 min at 4°C prior of adding into the streptavidin agarose. The reaction mixtures were incubated for 2 hours at 4°C. Agarose beads were then washed four times with binding buffer (50 mM Tris-HCl pH 7.5, 200 mM NaCl, 0.1% Triton X-100, 5% glycerol and 2 mM β-mercaptoethanol) and the samples of beads were analyzed by SDS-PAGE.

#### Alphafold prediction

To obtain the model of VPS29C and VPS29C containing Retromer, we applied the AlphaFold2 neural network of the open-source ColabFold pipeline. For each modelling experiment, ColabFold was excused using default settings where multiple sequence alignments were generated with MMseqs2. Five models were generated with the top 1 ranked model further relaxed using Amber. Structural alignments and molecular figures were generated using PyMOL and ChimeraX 1.6.1.

#### Phylogenetic Analysis of VPS29 Sequences

The full sequence and extended motif (AA 2-29) of Human VPS29C from UniProt (UniProt Consortium, 2019) (F8VXU5) was used as a query for initial BLAST searches (with default settings) against other metazoans. When BLASTing using only the extended motif, only one non-primate sequence was found: *<u>Bison bison bison</u>* (XP_010839672.1). To evaluate whether the amino-terminal extension might be present but unannotated in other animal genomes spanning the phylogenetic distance between primates and bison, we performed protein-to-genome searches using miniprot v0.12 (Li, 2023) against a representative selection of metazoans from the NCBI RefSeq database, using default settings, the –aln and –gff flags, and using human, bison and subsequently sloth VPS29C sequences as queries. These searches identified VPS29C amino-terminal extensions in range of additional placental mammals, but not in more distantly related lineages. The translated amino acid sequences were aligned using mafft v7.05 (L-INS-i) (Katoh et al., 2005) and visually inspected to find homologous motifs (Figure S1A). Multiple rounds of maximum likelihood tree inference were used to identify homologous sequences using IQTREE v2.2.5 (Minh et al., 2020), with the best fitting substitution model (-m MFP) and 10000 ultrafast bootstraps (-B 10000).

#### Proteomics

##### Experimental Design

GFP-IP experiments were performed as described above, and following the final wash the beads were processed for mass spectrometry to identify binding partners. All proteomic experiments were performed with isobaric tandem mass tagging (TMT) followed by LC-MS/MS quantitative mass spectrometry. The effects of TMT ratio suppression were minimised by pre-fractionation of the TMT-labelled pool and use of SPS-MS3-based acquisition to minimise ratio suppression due to co-isolation of peptides and, where possible, selecting the labelling set up to minimise any effects of channel bleed-through (Brenes et al., 2019).

##### TMT Labelling and High pH reversed-phase chromatography

GFP-IP samples were reduced (10mM TCEP, 55°C for 1h), alkylated (18.75mM iodoacetamide, room temperature for 30min.) and then digested from the beads with trypsin (2.5µg trypsin; 37°C, overnight). The resulting peptides were labelled with Tandem Mass Tag (TMTpro) sixteen plex reagents according to the manufacturer’s protocol (Thermo Fisher Scientific, Loughborough, LE11 5RG, UK) and the labelled samples pooled.

The pooled sample was evaporated to dryness, resuspended in 5% formic acid and then desalted using a SepPak cartridge according to the manufacturer’s instructions (Waters, Milford, Massachusetts, USA). Eluate from the SepPak cartridge was again evaporated to dryness and resuspended in buffer A (20 mM ammonium hydroxide, pH 10) prior to fractionation by high pH reversed-phase chromatography using an Ultimate 3000 liquid chromatography system (Thermo Scientific). In brief, the sample was loaded onto an XBridge BEH C18 Column (130Å, 3.5 µm, 2.1 mm X 150 mm, Waters, UK) in buffer A and peptides eluted with an increasing gradient of buffer B (20 mM Ammonium Hydroxide in acetonitrile, pH 10) from 0-95% over 60 minutes. The resulting fractions (concatenated into 5 in total) were evaporated to dryness and resuspended in 1% formic acid prior to analysis by nano-LC MSMS using an Orbitrap Fusion Tribrid mass spectrometer (Thermo Scientific).

##### Nano-LC Mass Spectrometry

High pH RP fractions were further fractionated using an Ultimate 3000 nano-LC system in line with an Orbitrap Fusion Tribrid mass spectrometer (Thermo Scientific). In brief, peptides in 1% (vol/vol) formic acid were injected onto an Acclaim PepMap C18 nano-trap column (Thermo Scientific). After washing with 0.5% (vol/vol) acetonitrile 0.1% (vol/vol) formic acid peptides were resolved on a 250 mm × 75 μm Acclaim PepMap C18 reverse phase analytical column (Thermo Scientific) over a 150 min organic gradient, using 7 gradient segments (1-6% solvent B over 1min., 6-15% B over 58min., 15-32%B over 58min., 32-40%B over 5min., 40-90%B over 1min., held at 90%B for 6min and then reduced to 1%B over 1min.) with a flow rate of 300 nl min^−1^. Solvent A was 0.1% formic acid and Solvent B was aqueous 80% acetonitrile in 0.1% formic acid. Peptides were ionized by nano-electrospray ionization at 2.0kV using a stainless-steel emitter with an internal diameter of 30 μm (Thermo Scientific) and a capillary temperature of 275°C.

All spectra were acquired using an Orbitrap Fusion Tribrid mass spectrometer controlled by Xcalibur 2.1 software (Thermo Scientific) and operated in data-dependent acquisition mode using an SPS-MS3 workflow. FTMS1 spectra were collected at a resolution of 120 000, with an automatic gain control (AGC) target of 400 000 and a max injection time of 100ms. Precursors were filtered with a minimum intensity of 5000, according to charge state (to include charge states 2-6) and with monoisotopic peak determination set to peptide. Previously interrogated precursors were excluded using a dynamic window (60s +/-10ppm). The MS2 precursors were isolated with a quadrupole isolation window of 1.2m/z. ITMS2 spectra were collected with an AGC target of 10 000, max injection time of 70ms and CID collision energy of 35%.

For FTMS3 analysis, the Orbitrap was operated at 30,000 resolution with an AGC target of 50 000 and a max injection time of 105ms. Precursors were fragmented by high energy collision dissociation (HCD) at a normalised collision energy of 55% to ensure maximal TMT reporter ion yield. Synchronous Precursor Selection (SPS) was enabled to include up to 10 MS2 fragment ions in the FTMS3 scan.

##### Data Analysis

The raw data files were processed and quantified using Proteome Discoverer software v2.4 (Thermo Scientific) and searched against the UniProt Human database (downloaded January 2022; 178486 sequences), an in-house ‘contaminants’ database and against the GFP sequence using the SEQUEST HT algorithm. Peptide precursor mass tolerance was set at 10ppm, and MS/MS tolerance was set at 0.6Da. Search criteria included oxidation of methionine (+15.995Da), acetylation of the protein N-terminus (+42.011Da) and methionine loss plus acetylation of the protein N-terminus (-89.03Da) as variable modifications and carbamidomethylation of cysteine (+57.021Da) and the addition of the TMTpro mass tag (+304.207Da) to peptide N-termini and lysine as fixed modifications. Searches were performed with full tryptic digestion and a maximum of 2 missed cleavages were allowed. The reverse database search option was enabled and all data was filtered to satisfy false discovery rate (FDR) of 5%.

#### Statistics

Further processing and statistical analysis of proteomics data was performed in the R statistical environment (version 4.2. PD2.4 protein grouping was retained, however, the master protein selection was improved with an in-house script which first searches Uniprot for the current status of all protein accessions and updates redirected or obsolete accessions. The script further takes the candidate master proteins for each group, and uses current uniprot review and annotation status to select the best annotated protein as master protein without loss of identification or quantification quality.

Protein abundances were normalised to GFP, and both raw and normalised abundances were then log_2_-transformed to bring them closer to a normal distribution prior to statistical analysis. Differential protein abundance was estimated using a Welch’s t-test (p-value). Volcano plots were plotted in GraphPad Prism 9 software (LaJolla, CA). Protein-protein interaction network analysis was performed using Metascape 3.5 (Zhou et al., 2019) and visualised using Cytoscape 3.3 software with the Enrichment Map plug-in (Merico et al., 2010). Dotplots were generated using ProHits-viz (Knight et al., 2017).

All statistical analysis was performed on data from a minimum of 3 independent experimental repeats. Unpaired t-tests were used to compare between the means of two groups, and one- or two-way ANOVA with Dunnett’s or Tukey’s multiple comparisons tests were used to compare the means of more than two groups. Graphs were prepared in GraphPad Prism 9. Individual datapoints represent independent experimental repeats. Graphs are plotted representing the mean value ± the standard error of the mean (SEM) for each experimental condition. *n* represents the number of independent experimental repeats. In all graphs, * = p < 0.05, ** = p < 0.01, *** = p < 0.001, **** = p < 0.0001.

## REFERENCES

Baños-Mateos S, Rojas AL, Hierro A. (2019). VPS29, a tweak tool of endosomal recycling. Curr Opin Cell Biol 59:81–87.

Bärlocher K, Hutter CAJ, Swart AL, Steiner B, Welin A, Hohl M, Letourneur F, Seeger MA, Hilbi H. (2017). Structural insights into Legionella RidL-VPS29 retromer subunit interaction reveal displacement of the regulator TBC1D5. Nat Commun 8, 1543.

Bartuzi P, Billadeau DD, Favier R, Rong S, Dekker D, Fedoseienko A, Fieten H, Wijers M, Levels JH, Huijkman N, et al. (2016). CCC- and WASH-mediated endosomal sorting of LDLR is required for normal clearance of circulating LDL. Nat Commun 7, 10961.

Boesch DJ, Singla A, Han Y, Kramer DA, Liu Q, Suzuki K, Juneja P, Zhao X, Long X, Medlyn MJ, et al. (2024). Structural organisation of the retriever-CCC endosomal recycling complex. Nat Struct Mol Biol 31, 910–924.

Brenes A, Hukelmann J, Bensaddek D, Lamond AI. (2019). Multibatch TMT reveals false positives, batch effects and missing values. Mol Cell Proteomics 18:1967–1980.

Bugarcic A, Zhe Y, Kerr MC, Griffin J, Collins BM, Teasdale RD. (2011). Vps26A and Vps26B subunits define distinct retromer complexes. Traffic 12, 1759–1773.

Bugarcic A, Vetter I, Chalmers S, Kinna G, Collins BM, Teasdale RD. (2015). Vps26B-retromer negatively regulates plasma membrane resensitization of PAR-2. Cell Biol Int 39, 1299–1306.

Butkovič R, Walker AP, Healy MD, McNally KE, Liu M, Kato K, Collins BM, Cullen PJ. (2024). Mechanism and regulation of cargo entry into the Commander recycling pathway. bioRxiv 2024.01.10.574988.

Carlton J, Bujny M, Peter BJ, Oorschot VM, Rutherford A, Mellor H, Klumperman J, McMahon HT, Cullen PJ. (2004). Sorting nexin-1 mediated tubular endosome-to-TGN transport through coincidence sensing of high-curvature and 3-phosphoinositides. Curr Biol 14, 1791–1800.

Chen KE, Healy MD, Collins BM. (2019). Towards a molecular understanding of endosomal trafficking by Retromer and Retriever. Traffic 20:465–478.

Chen KE, Guo Q, Hill TA, Cui Y, Kendall AK, Yang Z, Hall RJ, Healy MD, Sacharz J, Norwood SJ, et al. (2021). De novo macrocyclic peptides for inhibiting, stabilizing, and probing the function of the retromer endosomal trafficking complex. Sci Adv 7, eabg4007.

Chen K-E, Tillu VA, Gopaldass N, Chowdhury SR, Leneva N, Kovtun O, Ruan J, Guo Q, Ariotti N, Mayer A, Collins BM. (2024). Molecular basis for the assembly of the Vps5-Vps17 SNX-BAR proteins with Retromer. bioRxiv 2024.03.24.586500.

Clairfeuille T, Mas C, Chan AS, Yang Z, Tello-Lafoz M, Chandra M, Widagdo J, Kerr MC, Paul B, Merida I, Teasdale RD, Pavlos NJ, Anggono V, Collins BM. (2016). A molecular code for endosomal recycling of phosphorylated cargos by SNX27-retromer complex. Nat Struct Mol Biol 23, 921–932.

Collins BM, Skinner CF, Watson PJ, Seaman MN, Owen DJ. (2005). Vps29 has a phosphoesterase fold that acts as a protein interaction scaffold for retromer assembly. Nat Struct Mol Biol 12, 594–602.

Collins BM, Norwood SJ, Kerr MC, Mahony D, Seaman MN, Teasdale RD, Owen DJ. (2008). Structure of Vps26B and mapping of its interaction with the retromer protein complex. Traffic 9, 366–379.

Crawley-Snowdon H, Yang JC, Zaccai NR, Davis LJ, Wartosch L, Herman EK, Bright NA, Swarbrick JS, Collins BM, Jackson LP, et al. (2020). Mechanism and evolution of the Zn-fingernail required for interaction of VARP with VPS29. Nat Commun 11, 5031.

Cullen PJ, Steinberg F (2018). To degrade or not to degrade: mechanisms and significance of endocytic recycling. Nat Rev Mol Cell Biol 19, 679–696.

Dai J, Li J, Bos E, Porcionatto M, Premont RT, Bourgoin S, Peters PJ, Hsu VW. (2004). ACAP1 promotes endocytic recycling by recognizing sorting signals. Dev Cell 7, 771–776.

Damen E, Krieger E, Nielsen JE, Eygensteyn J, van Leeuwen JE. (2006). The human Vps29 retromer component is a metallo-phosphoesterase for a cation-independent mannose 6-phosphate receptor substrate peptide. Biochem J 398, 399–409.

Derivery E, Sousa C, Gautier JJ, Lombard B, Loew D, Gautreau A. (2009). The Arp2/3 activator WASH controls the fission of endosomes through a large multiprotein complex. Dev Cell 17, 712–723.

Freeman CL, Hesketh G, Seaman MN. (2014). RME-8 coordinates the activity of the WASH complex with the function of the retromer SNX dimer to control endosomal tubulation. J Cell Sci 127, 2053–2070.

Gallon M, Clairfeuille T, Steinberg F, Mas C, Ghai R, Sessions RB, Teasdale RD, Collins BM, Cullen PJ. (2014). A unique PDZ domain and arrestin-like fold interaction reveals mechanistic details of endocytic recycling by SNX27-retromer. Proc Natl Acad Sci USA 111, E3604–E3613.

Gilleron J, Zeigerer A (2023). Endosomal trafficking in metabolic homeostasis and diseases. Nat Rev Endocrinol 19, 28–45.

Giudice J, Xia Z, Wang ET, Scavuzzo MA, Ward AJ, Kalsotra A, Wang W, Wehrens XH, Burge CB, Li W, Cooper TA. (2014). Alternative splicing regulates vesicular trafficking genes in cardiomyocytes during postnatal heart development. Nat Commun 5:3603.

Gomez TS, Billadeau DD. (2009). A FAM21-containing WASH complex regulates retromer-dependent sorting. Dev Cell 17, 699–711.

Guo Q, Chen KE, Gimenez-Andres M, Jellett AP, Gao Y, Simonetti B, Liu M, Danson CM, Heesom KJ, Cullen PJ, Collins BM. (2024). Structural basis for coupling of the WASH subunit FAM21 with the endosomal SNX27-Retromer complex. Proceed Natl Acad Sci USA in press.

Gustavsson EK, Guella I, Trinh J, Szu-Tu C, Rajput A, Rajput AH, Steele JC, McKeown M, Jeon BS, Aasly JO, Farrer MJ. (2015). Genetic variability of the retromer cargo recognition complex in parkinsonism. Mov Disord 30:580–584.

Haft CR, de la Luz Sierra M, Bafford R, Lesniak MA, Barr VA, Taylor SI. (2000). Human orthologs of yeast vacuolar protein sorting proteins Vps26, 29, and 35: assembly into multimeric complexes. Mol Biol Cell 11, 4105–4116.

Harbour ME, Breusegem SYA, Antrobus R, Freeman C, Reid E, Seaman MNJ. (2010). The cargo-selective retromer complex is a recruiting hub for protein complexes that regulate endosomal tubule dynamics. J Cell Sci 123, 3703–3717.

Harbour ME, Breusegem SY, Seaman MN. (2012). Recruitment of the endosomal WASH complex is mediated by the extended ‘tail’ of Fam21 binding to the retromer protein Vps35. Biochem J 442, 209–220.

Harterink M, Port F, Lorenowicz MJ, McGough IJ, Silhankova M, Betist MC, van Weering JRT, van Heesbeen RGHP, Middelkoop TC, Basler K, Cullen PJ, Korswagen HC. (2011). A SNX3-dependent retromer pathway mediates retrograde transport of the Wnt sorting receptor Wntless and is required for Wnt secretion. Nat Cell Biol 13, 914–923.

Healy MD, McNally KE, Butkovič R, Chilton M, Kato K, Sacharz J, McConville C, Moody ERR, Shaw S, Planelles-Herrero VJ, et al. (2023). Structure of the endosomal Commander complex linked to Ritscher-Schinzel syndrome. Cell 186, 2219–2237.

Hesketh GG, Perez-Dorado I, Jackson LP, Wartosch L, Schafer IB, Gray SR, McCoy AJ, Zeldin OB, Garman EF, Harbour ME, et al. (2014). VARP is recruited on to endosomes by direct interaction with retromer, where together they function in export to the cell surface. Dev Cell 29, 591–606.

Hierro A, Rojas AL, Rojas R, Murthy N, Effantin G, Kajava AV, Steven AC, Bonifacino JS, Hurley JH. (2007). Functional architecture of the retromer cargo-recognition complex. Nature 449, 1063–1067.

Huotari J, Helenius A. (2011). Endosome maturation. EMBO J 30, 3481–3500.

Hsu JW, Bai M, Li K, Yang JS, Chu N, Cole PA, Eck MJ, Li J, Hsu VW. (2020). The protein kinase Akt acts as a coat adaptor in endocytic recycling. Nat Cell Biol 22, 927–933.

Jia D, Gomez TS, Billadeau DD, Rosen MK. (2012). Multiple repeat elements within the FAM21 tail link the WASH actin regulatory complex to the retromer. Mol Biol Cell 23, 2352–2361.

Jia D, Zhang JS, Li F, Wang J, Deng Z, White MA, Osborne DG, Phillips-Krawczak C, Gomez TS, Li H, et al. (2016). Structural and mechanistic insights into regulation of the retromer coat by TBC1d5. Nat Commun 7, 13305.

Jimenez-Orgaz A, Kvainickas A, Nagele H, Denner J, Eimer S, Dengjel J, Steinberg F. (2018). Control of RAB7 activity and localization through the retromer-TBC1D5 complex enables RAB7-dependent mitophagy. EMBO J 37, 235–254.

Katoh K, Kuma K, Toh H, Miyata T. (2005). MAFFT version 5: improvement in accuracy of multiple sequence alignment. Nucleic Acid Res 33:511–518.

Kendall AK, Xie B, Xu P, Wang J, Burcham R, Frazier MN, Binshtein E, Wei H, Graham TR, Nakagawa T, Jackson LP. (2020). Mammalian retromer is an adaptable scaffold for cargo sorting from endosomes. Structure 28, 393–405.

Kendall AK, Chandra M, Xie B, Wan W, Jackson LP. (2022). Improved mammalian retromer cryo-EM structures reveal a new assembly interface. J Biol Chem 298, 102523.

Kerr MC, Bennetts JS, Simpson F, Thomas EC, Flegg C, Gleeson PA, Wicking C, Teasdale RD. (2005). A novel mammalian retromer component, Vps26B. Traffic 6, 991–1001.

Kim E, Lee Y, Lee HJ, Kim JS, Song BS, Huh JW, Lee SR, Kim SU, Kim SH, Hong Y, Shim I, Chang KT. (2010). Implication of mouse Vps26b-Vps29-Vps35 retromer complex in sortilin trafficking. Biochem Biophys Res Commun 403, 167–171.

Klumperman J, Raposo G. (2014). The complex ultrastructure of the endolysosomal system. Cold Spring Harb Perspect Biol 6, a016857.

Knight JDR, Choi H, Gupta GD, Pelletier L, Raught B, Nesvizhskii AI, Gingras A-C. (2017). ProHits-viz: a suite of web tools for visualising interaction proteomics data. Nat Methods 14:645–646.

Kovtun O, Leneva N, Bykov YS, Ariotti N, Teasdale RD, Schaffer M, Engel BD, Owen DJ, Briggs JAG, Collins BM. (2018). Structure of the membrane-assembled retromer coat determined by cryo-electron tomography. Nature 561, 561–564.

Kvainickas A, Nagele H, Qi W, Dokladal L, Jimenez-Orgaz A, Stehl L, Gangurde D, Zhao Q, Hu Z, Dengjel J, et al. (2019). Retromer and TBC1D5 maintain late endosomal RAB7 domains to enable amino acid-induced mTORC1 signaling. J Cell Biol 218, 3019–3038.

Lauffer BE, Melero C, Temkin P, Lei C, Hong W, Kortemme T, von Zastrow M. (2010). SNX27 mediates PDZ-directed sorting from endosomes to the plasma membrane. J Cell Biol 190, 565–574.

Laulumaa S, Kumpula E-P, Huiskonen JT, Varjosalo M. (2024). Structure and interactions of the endogenous human Commander complex. Nat Struct Mol Biol 31, 925–938.

Leneva N, Kovtun O, Morado DR, Briggs JAG, Owen DJ. (2021). Architecture and mechanism of metazoan retromer:SNX3 tubular coat assembly. Sci Adv 7, eabf8598.

Li H. (2023). Protein-to-genome alignment with miniprot. Bioinformatics 39:btad014.

Li J, Peters PJ, Bai M, Dai J, Bos E, Kirchhausen T, Kandror KV, Hsu VW. (2007). An ACAP1-containing clathrin coat complex for endocytic recycling. J Cell Biol 178, 453–464.

Lucas M, Gershlick DC, Vidaurrazaga A, Rojas AL, Bonifacino JS, Hierro A. (2016). Structural mechanism for cargo recognition by the retromer complex. Cell 167, 1623–1635.

Mallam AL, Marcotte EM. (2017). Systems-wide studies uncover Commander, a multiprotein complex essential to human development. Cell Syst 4, 483–494.

Merico D, Isserlin R, Stueker O, Emili A, Bader GD. (2010). Enrichment map: a network-based method for gene-set enrichment visualisation and interpretation. PLoS One 5:e13984.

McGough IJ, Steinberg F, Jia D, Barbuti PA, McMillan KJ, Heesom KJ, Whone AL, Caldwell MA, Billadeau DD, Rosen MK, Cullen PJ. (2014a). Retromer binding to FAM21 and the WASH complex is perturbed by the Parkinson disease-linked VPS35(D620N) mutation. Curr Biol 24, 1678.

McGough IJ, Steinberg F, Gallon M, Yatsu A, Ohbayashi N, Heesom KJ, Fukuda M, Cullen PJ. (2014b). Identification of molecular heterogeneity in SNX27-retromer-mediated endosome-to-plasma membrane recycling. J Cell Sci 127, 4940–4953.

McGough IJ, de Groot REA, Jellett AP, Betist MC, Varandas KC, Danson CM, Heesom KJ, Korswagen HC, Cullen PJ. (2018). SNX3-retromer requires an evolutionary conserved MON2:DOPEY2:ATP9A complex to mediated Wntless sorting and Wnt secretion. Nat Commun 9, 3737.

McMillan KJ, Gallon M, Jellett AP, Clairfeuille T, Tilley FC, McGough IJ, Danson CM, Heesom KJ, Wilkinson KA, Collins BM, Cullen, PJ (2016). Atypical parkinsonism-associated retromer mutant alters endosomal sorting of specific cargo proteins. J Cell Biol 214, 389–399.

McMillan KJ, Korswagen HC, Cullen PJ (2017). The emerging role of retromer in neuroprotection. Curr Opin Cell Biol 47, 72–82.

McNally KE, Faulkner R, Steinberg F, Gallon M, Ghai R, Pim D, Langton P, Pearson N, Danson CM, Nägele H, et al. (2017). Retriever is a multiprotein complex for retromer-independent endosomal cargo recycling. Nat Cell Biol 19, 1214–1225.

McNally KE, Cullen PJ. (2018). Endosomal retrieval of cargo: retromer is not alone. Trends Cell Biol 28, 807–822.

Migliano SM, Wenzel EM, Stenmark H. (2022). Biophysical and molecular mechanisms of ESCRT function, and their implications for disease. Curr Opin Cell Biol 75, 102062.

Minh BQ, Schmidt HA, Chernomor O, Schrempf D, Woodhams MD, von Haeseler A, Lanfear R. (2020). IQ-TREE 2: New models and efficient methods for phylogenetic inference in the genomic era. Mol Biol Evol 37:1530–1534.

Moulay G, Laine J, Lemaitre M, Nakamori M, Nishino I, Caillol G, Mamchaoui K, Julien L, Dingli F, Loew D, et al. (2020). Alternative splicing of clathrin heavy chain contributes to the switch from coated pits to plaques. J Cell Biol 219:e201912061.

Norris A, Grant BD. (2020). Endosomal microdomains: formation and function. Curr Opin Cell Biol 65, 86–95.

Norwood SJ, Shaw DJ, Cowieson NP, Owen DJ, Teasdale RD, Collins BM. (2011). Assembly and solution structure of the core retromer protein complex. Traffic 12, 56–71.

Phillips-Krawczak CA, Singla A, Starokadomskyy P, Deng Z, Osborne DG, Li H, Dick CJ, Gomes TS, Koenecke M, Zhang JS, Dai H, Sifuentes-Dominguez LF, Geng LN, Kaufmann SH, Hein MY, Wallis M, McGaughran J, Gecz J, van de Sluis B, Billadea DD, Burstein E. (2015). COMMD1 is linked to the WASH complex and regulates endosomal trafficking of the copper transporter ATP7A. Mol Biol Cell 26, 91–103.

Puthenveedu MA, Lauffer B, Temkin P, Vistein R, Carlton P, Thorn K, Taunton J, Weiner OD, Parton RG, von Zastrow M. (2010). Sequence-dependent sorting of recycling proteins by actin-stabilised endosomal microdomains. Cell 143, 761–773.

Romano-Moreno M, Rojas AL, Williamson CD, Gershlick DC, Lucas M, Isupov MN, Bonifacino JS, Machner MP, Hierro A. (2017). Molecular mechanism for the subversion of the retromer coat by the Legionella effector RidL. Proc Natl Acad Sci USA 114, E11151–E11160.

Romano-Moreno M, Astorga-Simon EN, Rojas AL, Hierro A. (2024). Retromer-mediated recruitment of the WASH complex involves discrete interactons between VPS35, VPS29, and FAM21. Protein Sci 33, e4980.

Rowland A, Chitwood P, Phillips M, Voeltz G. (2014). ER contact sites define the position and timing of endosome fission. Cell 159, 1027–1041.

Seaman MN, McCaffery JM, Emr SD. (1998). A membrane coat complex essential for endosome-to-Golgi retrograde transport in yeast. J Cell Biol 142, 665–681.

Seaman MN, Harbour ME, Tattersall D, Read E, Bright N. (2009). Membrane recruitment of the cargo-selective retromer subcomplex is catalysed by the small GTPase Rab7 and inhibited by the Rab-GAP TBC1D5. J Cell Sci 122, 2371–2382.

Seaman MNJ, Mukadam AS, Breusegem SY. (2018). Inhibition of TBC1D5 activates Rab7a and can enhance the function of the retromer cargo-selective complex. J Cell Sci 131, jcs217398.

Shortill SP, Frier MS, Davey M, Conibear E. (2024). N-terminal signals in the SNX-BAR paralogs Vps5 and Vin1 guide endosomal coat complex formation. Mol Biol Cell 35, ar76.

Sigismund S, Lanzetti L, Scita G, Di Fiore PP. (2021). Endocytosis in the context-dependent regulation of individual and collective cell properties. Nat Rev Mol Cell Biol 22, 625–643.

Simoes S, Guo J, Buitrago L, Qureshi YH, Feng X, Kothiya M, Cortes E, Patel V, Kannan S, Kim YH, et al. (2021). Alzheimer’s vulnerable brain region relies on a distinct retromer core dedicated to endosomal recycling. Cell Rep 37, 110182.

Simonetti B, Paul B, Chaudhari K, Weeratunga S, Steinberg F, Gorla M, Heesom KJ, Bashaw GJ, Collins BM, Cullen PJ. (2019). Molecular identification of a BAR domain-containing coat complex for endosomal recycling of transmembrane proteins. Nat Cell Biol 21, 1219–1233.

Simonetti B, Guo Q, Gimenez-Andres M, Chen KE, Moody ERR, Evans AJ, Chandra M, Danson CM, Williams TA, Collins BM, Cullen PJ. (2022). SNX27-retromer directly binds ESCPE-1 to transfer cargo proteins during endosomal recycling. PLoS Biol 20, e3001601.

Small SA, Simoes-Spassov S, Mayeux R, Petsko GA. (2017). Endosomal traffic jams represent a pathogenic hub and therapeutic target in Alzheimer’s disease. Trends Neurosci 40, 592–602.

Steinberg F, Gallon M, Winfield M, Thomas EC, Bell AJ, Heesom KJ, Tavare JM, Cullen PJ. (2013). A global analysis of SNX27-retromer assembly and cargo specificity reveals a function in glucose and metal ion transport. Nat Cell Biol 15, 461–471.

Strochlic TI, Setty TG, Sitaram A, Burd CG. (2007). Grd19/Snx3p functions as a cargo-specific adapter for retromer-dependent endocytic recycling. J Cell Biol 177, 115–125.

Sugatha J, Priya A, Raj P, Jaimon E, Swaminathan U, Jose A, Pucadyil TJ, Datta S. (2023). Insights into cargo sorting by SNX32 and its role in neurite outgrowth. eLife 12:e84396.

Temkin P, Lauffer B, Jager S, Cimermancic P, Krogan NJ, von Zastrow M. (2011). SNX27 mediated retromer tubule entry and endosome-to-plasma membrane trafficking of signalling receptors. Nat Cell Biol 13, 715–721.

Traer CJ, Rutherford AC, Palmer KJ, Wassmer T, Oakley J, Attar N, Carlton JG, Kremerskothen J, Stephens DJ, Cullen PJ. (2007). SNX4 coordinates endosomal sorting of TfnR with dynein-mediated transport into the endocytic recycling compartment. Nat Cell Biol 9, 1370–1380.

UniProt Consortium. (2019). UniProt: a worldwide hub of protein knowledge. Nucleic Acids Res 47:D506–D515.

van Kerkhof P, Lee J, McCormick L, Tetrault E, Lu W, Schoenfish M, Oorschot V, Strous GJ, Klumperman J, Bu G. (2005). Sorting nexin 17 facilitates LRP recycling in the early endosome. EMBO J 24, 2851–2861.

van Weering JR, Sessions RB, Traer CJ, Kloer DP, Bhatia VK, Stamou D, Carlsson SR, Hurley JH, Cullen PJ. (2012). Molecular basis for SNX-BAR-mediated assembly of distinct endosomal sorting tubules. EMBO J 31, 4466–4480.

Varandas KC, Irannejad R, von Zastrow M. (2016). Retromer endosome exit domains serve multiple trafficking destinations and regulate local G protein activation by GPCRs. Curr Biol 26, 3129–3142.

Wang D, Guo M, Liang Z, Fan J, Zhu Z, Zang J, Zhu Z, Li X, Teng M, Niu L, Dong Y, Liu P. (2005). Crystal structure of human vacuolar protein sorting protein 29 reveals a phosphodisesterase/nuclease-like fold and two protein-protein interactions sites. J Biol Chem 280, 22962–22967.

Wang H, Qi W, Zou C, Xie Z, Zhang M, Naito MG, Mifflin L, Liu Z, Najafov A, Pan H, Shan B, Li Y, Zhu ZJ, Yuan J. (2021). NEK1-mediated retromer trafficking promotes blood-brain barrier integrity by regulating glucose metabolism and RIPK1 activation. Nat Commun 12, 4826.

Yao J, Yang F, Sun X, Wang S, Gan N, Liu Q, Liu D, Zhang X, Niu D, Wei Y, et al. (2018). Mechanism of inhibition of retromer transport by the bacterial effector RidL. Proc Natl Acad Sci USA 115, E1446–E1454.

Yong X, Zhao L, Deng W, Sun H, Zhou X, Mao L, Hu W, Shen X, Sun Q, Billadeau DD, Xue Y, Jia D. (2020). Mechanism of cargo recognition by retromer-linked SNX-BAR proteins. PLoS Biol 18, e3000631.

Yong X, Zhao L, Hu W, Sun Q, Ham H, Liu Z, Ren J, Zhang Z, Zhou Y, Yang Q, et al. (2021). SNX27-FERM-SNX1 complex structure rationalizes divergent trafficking pathways by SNX17 and SNX27. Proc Natl Acad Sci USA 118, e2105510118.

Young JE, Holstege H, Andersen OM, Petsko GA, Small SA. (2023). On the causal role of retromer-dependent endosomal recycling in Alzheimer’s disease. Nat Cell Biol 25, 1394–1397.

Zhou Y, Zhou B, Pache L, Chang M, Khodabakshi AH, Tanaseichuk O, Benner C, Chanda SK. (2019). Metascape provides a biologist-oriented resource for the analysis of systems-level datasets. Nat Commun 10:1523.

