## Supplementary Table 6 for "Identification of a VPS29 isoform with restricted association to Retriever and Retromer accessory proteins through auto-inhibition"

**Supplementary Table 6. Thermodynamic parameters for the binding of Vps29C, Vps35L and cyclic peptides by ITC.**

|  | ***K_d_***  **(μM)** | **Δ*H* (kcal/mol)** | **Δ*G* (kcal/mol)** | ***-T*Δ*S* (kcal/mol)** |
| --- | --- | --- | --- | --- |
| **GST-VPS29C** |  |  |  |  |
| VPS35L 16-38 | >100 | -9.54 ± 2.50 | -5.26 ± 0.15 | 4.26 ± 2.04 |
| VPS35L 28-37 | No binding detected | | | |
| RT-D1 L7E | No binding detected | | | |
| RT-D3 | 13.40 ± 1.67 | -2.02 ± 0.71 | -6.65 ± 0.07 | -5.06 ± 1.44 |
| **GST-VPS29B** |  |  |  |  |
| RT-D3 | 0.014 ± 0.001 | -10.40 ± 0.42 | -2.81 ± 0.01 | -1.72 ± 1.53 |
